# Full-Atom MPNN Based Redesign of Plant Dehydrogenase Enables Thermostability Enhancement Without Loss of Stereoselectivity

**DOI:** 10.64898/2026.04.20.719482

**Authors:** Bruno Di Geronimo, Jasmin Zuson, Ana Udženija, Andrea M. Chánique, Robert Kourist, Shina Caroline Lynn Kamerlin

**Author notes:** Corresponding author email addresses. These authors contributed equally (Bruno Di Geronimo, Jasmin Zuson). Author current address: Kura Biotech, Av. Gramado 1410 interior, Parcela 20, Puerto Varas 5550000, Chile.

## Abstract

Protein stabilization is a “Holy Grail” of biocatalysis, and stability design is an area of intense research interest. While it is increasingly feasible to effectively increase enzyme thermostability, optimization without compromising activity or selectivity remains a significant challenge. Here, we use full-atom protein sequence design with sidechain conditioning (FAMPNN) to engineer thermostable variants of the borneol dehydrogenase from *Salvia rosmarinus* (*Sr*BDH1), an enzyme from a family where unselective enzymes dominate, and selectivity is determined by dynamical considerations. By combining FAMPNN design with residue conservation analysis and avoiding active site residues, we were able to computationally design *Sr*BDH1 variants with up to 10 °C enhanced thermostability and strongly increased half-life time at elevated temperature, while retaining selectivity towards (+)-borneol. This design framework, integrating *de novo* and physics-based protein design tools, demonstrates that stability can be enhanced without disrupting functionally relevant dynamics, providing a route to engineer robust and selective biocatalysts.

Graphical Abstract

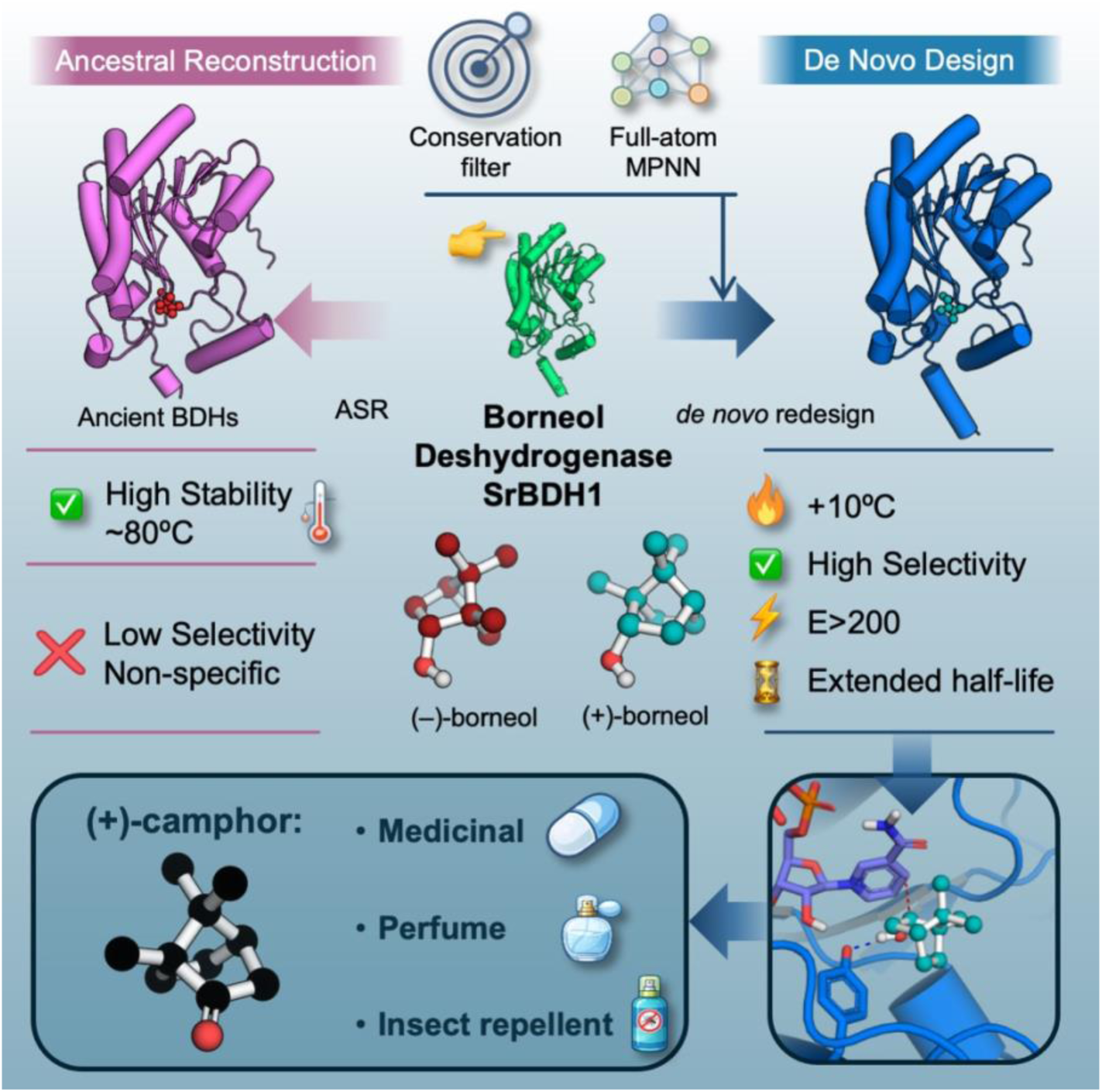

## Introduction

Protein stability engineering is considered a “Holy Grail” of biocatalysis, as it is essential for allowing enzymes to operate under harsh industrial conditions, such as high temperature and organic solvents.^1,2^ While an abundance of experimental and computational approaches exist to engineer stable proteins more broadly,^1–3^ a main challenge to be overcome is the function-stability tradeoff.^3–15^ Specifically, most natural proteins are only marginally stable,^3,16–18^ and mutations that improve protein function (for instance catalytic activity or ligand binding) often do so at the cost of protein stability. Conversely, attempts to improve stability can compromise catalytic function or enzyme selectivity.^3–15^

A number of strategies have been developed to address this issue. Among them, directed evolution is a powerful tool to find “compensatory” mutations – changes elsewhere in the protein scaffold that fix folding issues created by new functional mutations.^19–25^ Studies utilizing classic error-prone PCR (epPCR) report increased thermal stability after screening a minimum of 19,000 clones in one round of directed evolution.^26^ More complex combined approaches, aim to minimize the screening effort of the generated libraries by design, but still screen 7 to 8 thousand clones per round.^27^ However, achieving this is often only possible over multiple rounds of laboratory evolution, which carries both monetary and human time costs, and is likely to hit optimization plateaus.^28–32^ Another alternative is the use of ancestral sequence reconstruction (ASR),^33–35^ which tends to predict proteins with substantively higher thermostability,^36–40^ although the origins of this (reproduction of early earth conditions *vs.* algorithmic bias) remain controversial.^36,41,42^ However, while ASR is an effective tool to quickly obtain enhanced stability, the predicted variants can have solubility and expression challenges, or other issues such as for instance lower capacity to bind an internal cofactor.

More recently, there has been substantive interest in using pure phylogenetic,^39,43,44^ hybrid phylogenetic/physics-based ^44,45^ and machine learning-based approaches such as RFDiffusion,^46,47^ ProteinMPNN,^48^ ThermoMPNN,^49^ Evo-Velocity,^50^ and ESM-2/ESM-Fold,^51^ among other examples, to predict stable and functional protein scaffolds *de novo*. Recently, we used ASR to generate the ancestors of a heterogeneous family of plant borneol dehydrogenases (BDHs) that are associated to the oxidation of (+)-1-borneol to camphor.^52^ While most of these short-chain reductases accept both enantiomers of 1-borneol,^53–55^ the enzymes from rosemary and sage have outstanding enantioselectivity in the kinetic resolution of this diborane type monoterpenol.^56^ Not surprisingly, given function-stability tradeoffs,^3–15^ most ASR characterized ancestors were no longer enantioselective, with the exception of the direct ancestors of the enzymes from rosemary and sage.^52^ Similar to previously characterized ASR-generated proteins,^57,58^ shifts in selectivity across the transition from predicted ancestral to extant enzymes could be associated to perturbations in conformational dynamics.

Although ASR has been to some extent a “gold-standard” for easy acquisition of thermostability,^36–39^ it is clearly not without functional cost. Similarly to ASR, *de novo* designed proteins have also been exhibited to be highly thermostable.^59–63^ In the current work, we explore whether ML-based approaches can be used to obtain thermostability *without* compromising activity and stereoselectivity in *de novo* designed enzymes, using FAMPNN as a model approach, based on prior successes of this approach in enhancing protein thermostability.^64^ In addition, the last two decades have shown that focus of saturation mutagenesis into the active site is a powerful method to alter the selectivity of an enzyme.^65–69^ In many cases, variation of first shell amino acids is sufficient for such an improvement, and it is not necessary to address the periphery.^66^ In this work, we hypothesized that excluding active site amino acids from the redesign strategy will protect the active site, and retain selectivity. To test this hypothesis, we generated a set of 8 BDH variants by redesign of peripheral regions of the BDH from *Salvia rosmarinus* (*Sr*BDH1^54,70^), while conserving residues that are supposed to participate in the catalytic machinery and the active-site hydrophobic pockets.

Using this approach, coupled with experimental validation of computational designs, we demonstrate that we were able to obtain a variant with a 10°C higher unfolding temperature, and with minimal trade-off in activity (∼65% of the specific activity retained) and, crucially, while retaining selectivity, unlike in the ASR-based variants predicted in prior work.^52^ This is significant given that maintaining selectivity in *Sr*BDH1 appears to be particularly challenging.^52^ Finally, computational characterization of both the computationally redesigned sequences and the ancestral proteins from our prior ASR study^52^ indicated that thermostability and stereoselectivity are governed by distinct and partially decoupled structural features, with selectivity primarily emerging from the dynamic stabilization of catalytically competent conformations rather than static active-site geometry. While BDH is itself an industrially important enzyme for the synthesis of borane-type monoterpenoids,^52–54,71,72^ our study also highlights the power of strategic exploitation of ML-based approaches for protein stability design without compromising enzyme selectivity, which has major applications for biocatalysis and provides a possible avenue also for the stabilization of engineered enzyme variants having artificially improved stereoselectivity.

## Results and Discussion

### Sequence Generation and Thermostability Predictions

ConSurf analysis^73,74^ was performed on the borneol dehydrogenase of *Salvia rosmarinus* (*Sr*BDH1, PDB ID: 6ZYZ^54^), providing us with residue conservation cutoffs,^75^ normalized from 1 to 9, from more variable to highly conserved positions (**Figures 1A** and **S1**). We then applied two different thresholds (residue conservation cutoffs of ≤ 4 and ≤ 5), and then performed full-atom protein sequence design with sidechain conditioning using FAMPNN (Full-Atom MPNN), an inverse folding model that co-designs sequence and sidechains.^64^ Taking sidechains into account is important, as while other more specialized options such as ThermoMPNN^49^ also exist, our goal is not just to enhance thermostability, but to do so without compromising selectivity, which in turn is determined by sidechain conformations and dynamics.^76–80^ A further key advantage of FAMPNN^64^ is that it outputs *de novo Sr*BDH1 designs as fully resolved all-atom Protein Data Bank (PDB)^81^ structures, ensuring intrinsic structural self-consistency, and eliminating the need for post-hoc side-chain repacking. This also makes it easy to evaluate and rank FAMPNN^64^ output by feeding it into PyRosetta^82^ to compute ΔΔ*G* values.

**Figure 1.**
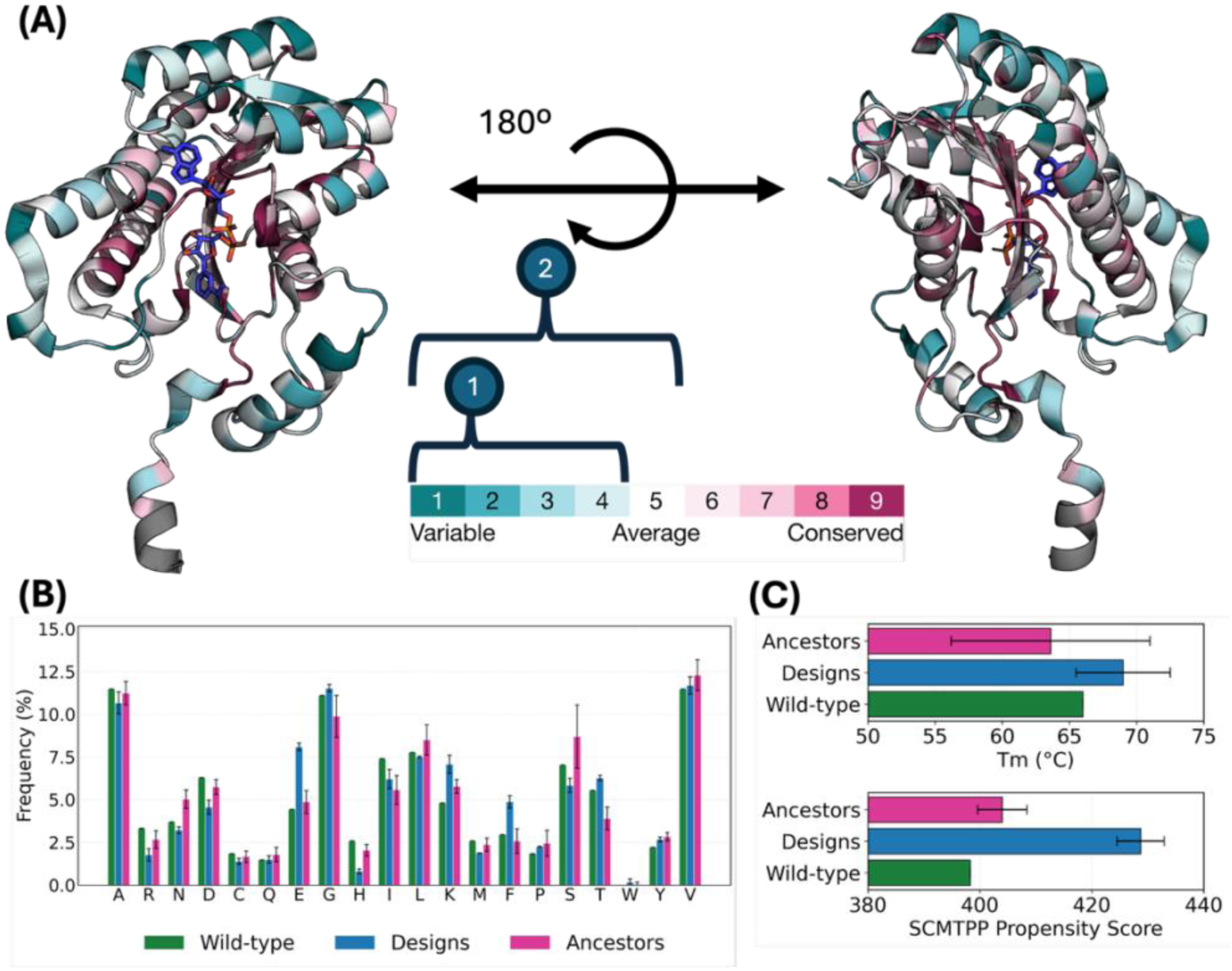
(**A**) Borneol dehydrogenase (BDH) residue conservation, performed using ConSurf,^73,74^ with the BDH from *Salvia rosmarinus* (*Sr*BDH1)^54^ used as a reference. Chain A from the *Sr*BDH1 structure (PDB ID: 6ZYZ^54^) is shown in cartoon representation, and colored by a ConSurf^73,74^ conservation score (1–9; cyan = variable → magenta = conserved). NAD⁺ is shown as blue sticks. Two orientations of this enzyme (left/right) are shown to highlight a conserved cluster of residues around the active-site pocket, adjacent to the NAD⁺ cofactor. (**B**) Amino acid composition of wild-type *Sr*BDH1 (green), designed sequences (blue) and ancestral sequences (magenta), performed using the ExPASy webserver.^83^ (**C**) Experimental thermostability mean values, together with SCMTPP^84^ (“Scoring Card Method Thermophilic Protein Predictor”)-derived propensity score predictions and standard deviations, for the wild-type system (green), designed *Sr*BDH1 variants (blue), and ancestral sequences (magenta).

Prior work has indicated a number of sequence and structural factors being important indicators of thermostability. These include: (1) hydrophobic content of the sequence,^85–88^ which can be measured using the Kyte-Doolittle scale,^89^ either as % hydrophobic content of the sequence, or through Grand Average of Hydropathicity scores (GRAVY index, where the GRAVY score is simply the sum hydropathy values of all amino acids divided by the total number of amino acids). (2) Related to this, statistical analysis suggests that globular proteins from thermophilic organisms have a significantly higher aliphatic index (a measure of the volume occupied by their aliphatic side chains) than their mesophilic counterparts.^90^ (3) Analysis of sequence composition has suggested that thermostable proteins show an increased frequency of Arg and Tyr side chains, and a reduced frequency of Cys and Ser side chains, compared to their mesophilic counterparts^91^ (in the case of hyperthermostable archaeal protein tyrosine phosphatases, as also observed a higher frequency of Glu and Leu residues^92^). (4) Finally, from a structural perspective, thermostable proteins have both a larger fraction of their residues in an α-helical conformation and avoid Pro to a larger extent in those α-helices,^91^ they have a higher amount of salt bridges and hydrogen bonds compared to mesophilic counterparts,^91^ and maintain a higher percentage of their residues engaged in hydrophobic contacts (in line with expected increased hydrophobic content).^92^

For sequence-based putative thermostability analysis, we compared two sets of sequences to that of wild-type *Sr*BDH1: (1) FAMPNN designs^64^ selecting the top 4 sequences based on Rosetta Energy Unit (REU) values^93^ from PyRosetta,^82^ with two design batches, *i.e.*, designs at residue positions with ConSurf^73,74^ residue conservation cutoffs of ≤ 4 and ≤ 5, and (2) a series of ASR predictions from our prior work.^52^ The sequences being considered are provided in the **Supplementary Information**, and the corresponding analysis is provided in **Figure 1**, with the corresponding raw data provided in the **Supplementary Table**). Amino acid composition, instability indices, aliphatic indices, and GRAVY scores were all calculated using the ProtParam tool on the Expasy webserver.^83^ Protein thermostability was predicted from sequence based on propensity scores calculated using the SCMTPP webserver, a sequence-based thermophilic protein predictor (TPP) that uses a scoring card method (SCM).^84^

Our sequence-based analysis indicates, most importantly, that both FAMPNN^64^ and ASR sequences have increased SCMTPP propensity scores compared to wild-type *Sr*BDH1, with the FAMPNN^64^ designs having on average even higher propensity scores than the ASR sequences, suggestive of the highest thermostability (398.3 for wild-type *Sr*BDH1, 404.0 ± 4.4 for the ASR sequences, and 428.7 ± 4.2 for the FAMPNN^64^ designs). In addition, the predicted sequences all also show lower GRAVY scores than the experimental designs, slightly lower instability indices, and similar aliphatic indices. Finally, in terms of sequence composition, curiously, we see a reduced frequency of Arg side chains and a very modest increase in frequency of Tyr side chains in the FAMPNN^64^ designs compared to wild-type, as well as a reduced frequency of Cys and Ser side chains compared to both wild-type and the ASR constructs, hinting at greater thermostability.^91^ Interestingly, we also see an increased frequency of Glu, and Phe residues in the designed sequences, compared to either wild-type or ASR sequences. Finally, despite their highlighted importance in prior work,^85–88,90^ we do not see meaningful trends in either aliphatic indices or GRAVY scores, that would distinguish between wild-type *Sr*BDH1 and the FAMPNN^64^ or ASR sequences.^52^

Following from this, we structurally and dynamically profiled wild-type *Sr*BDH1 and the *de novo* designed *Sr*BDH1 constructs, as well as three ancestral sequences from our prior work^52^ for reference, based on molecular dynamics simulations performed as described in the **Materials and Methods**. The three selected ancestral sequences, N32, N5 and N6 have *T*_m_ values ranging from 61-77 °C, see **Section “Experimental Characterization and Validation”**). Structural stability was evaluated by monitoring non-covalent interactions including hydrogen bonds, salt bridges, and hydrophobic interactions (**Figure 2**). Additionally, solvent accessible surface area (SASA) and root mean square fluctuations (RMSF) of backbone C_α_-atoms were calculated (**Figures S2** to **S4**, and **Table S1**).

**Figure 2.**
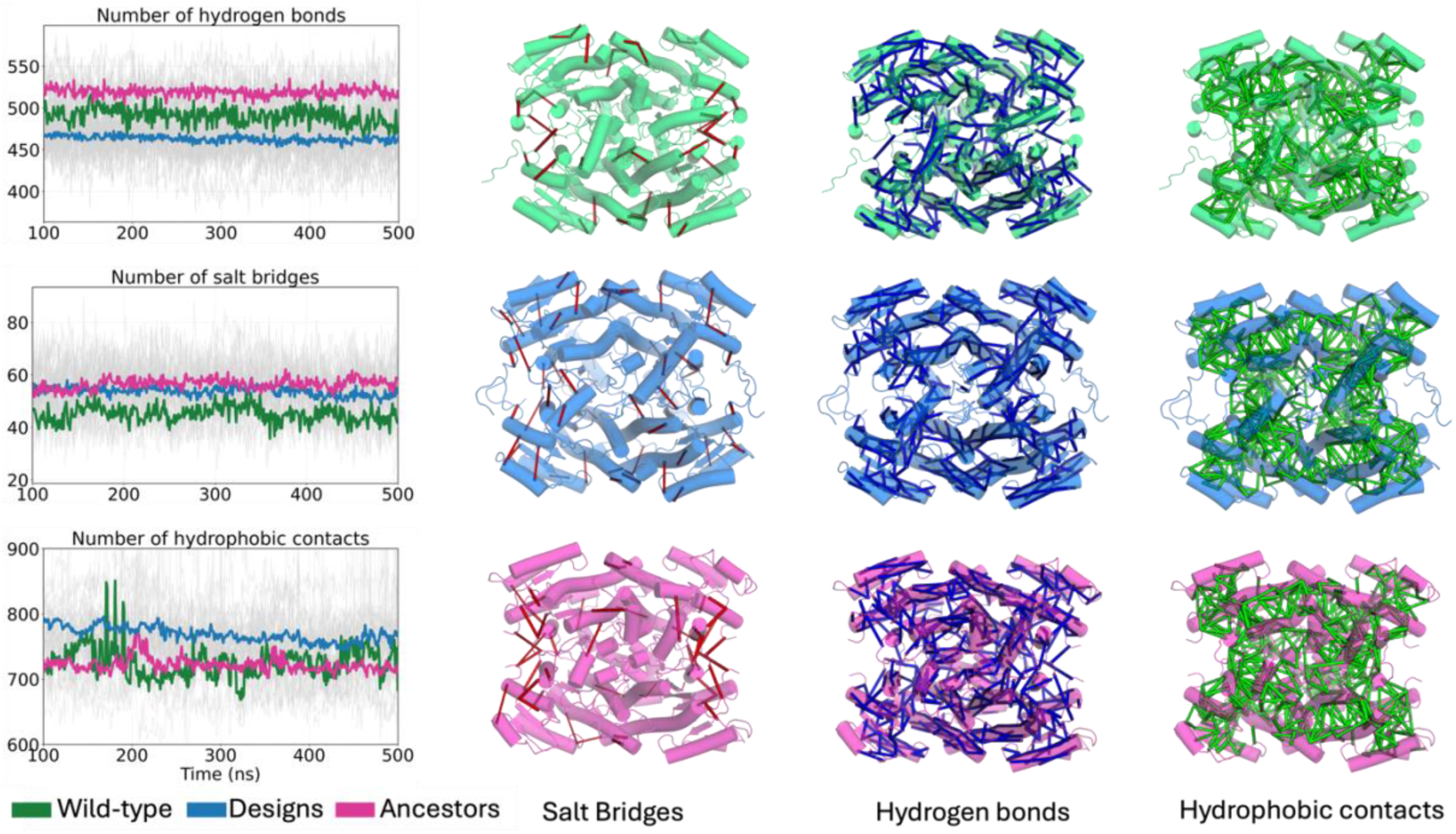
Computational analysis of non-covalent interaction networks associated with *Sr*BDH1 stability across wild-type, designed variants, and ancestral systems. Data are derived from the last 400 ns of 3 x 500 ns conventional MD simulations. Hydrogen bonds were identified using CPPTRAJ,^98^ (and salt bridges and hydrophobic contacts were quantified using MDAnalysis,^99,100^ with mean values for each system shown in color. The cartoon representations highlight the distribution of hydrogen bonds, salt bridges, and hydrophobic interactions mapped onto representative protein conformations, illustrating differences in interaction networks between wild-type (green), designed variants (blue), and ancestral proteins (magenta).

Analysis of non-covalent interactions during our simulations (**Figure 2**) reveals clear differences in interaction propensity between wild-type *Sr*BDH1, the new designed variants, and the ancestral *Sr*BDH1 variants from our prior work.^52^ The ancestral enzymes simulated display a higher number of hydrogen bonds compared to both the wild-type enzyme and designed variants, suggesting increased internal stabilization through polar interactions. In contrast, the designed variants exhibit an increased number of salt bridges, consistent with enhanced electrostatic stabilization, which has been suggested to be associated with increased protein thermostability.^94–97^ Moreover, hydrophobic contacts are more prevalent in the designed systems compared to both wild-type and ancestral variants, reflecting a redistribution of stabilizing interactions within the protein scaffold, likely driven by the increased abundance of phenylalanine residues in the designs.

Although these differences may appear moderate at the global level, they arise from specific residue substitutions that reorganize interaction networks across the protein scaffold. For example, comparison between the wild-type and design 01-080 (**Figure S5**) reveals the introduction of new salt bridges through the A27R, R31E, Q46R, and L49E as well as N60D, V135K, V135R, and S180K substitutions. In addition, hydrophobic interactions are enhanced by substitutions such as H85W, L122F, H158F and V176F (**Figures S5**), contributing to a more tightly packed and stabilized protein scaffold. Monitoring of the solvent accessible surface area (SASA) values (Å^2^) across variants, including both total and hydrophobic contributions, shows a consistent decrease in surface exposure relative to the wild-type enzyme for both designs and ancestral proteins (**Figures S2** and **Table S1**). This, in turn, is likely to contribute to enhanced compactness and structural robustness of the designs, potentially reinforced by an increased abundance of charged residues and the formation of stabilizing salt-bridges at the protein surface. Indeed, analysis of non-covalent interactions (**Figure S2B**) confirms that most FAMPNN designs^64^ show a reduction in total hydrogen bonds but an increase in salt bridge interactions over the course of the last 400 ns of simulation time (3 trajectories per system), with particularly pronounced increases observed for designs 02-037, 02-389, and 02-911, which reach mean values of approximately 62 salt bridges per system. In contrast, designs 01-037 and 01-173 have non-covalent interaction totals that are comparable to wild-type *Sr*BDH1. Further, while the total number of hydrogen bonds is generally reduced in the designs relative to the wild type, variants 01-080, 01-099, and 02-037 retain relatively high hydrogen bond counts throughout the simulations.

Following from this, we also dynamically profiled wild-type *Sr*BDH1 designed variants, and ancestors considering C_α_-atom RMSF across our MD simulations (**Figure 3**). We note that the relationship between protein thermostability and flexibility is complex,^101^ as are the structural features that correlated with protein thermal stability.^84,91^ The general view of protein flexibility is that protein thermostability is linked to increased scaffold rigidity.^101–107^ However, several studies have demonstrated that this is not necessarily the case, and that flexibility and thermal stability can be decoupled.^101,108–111^ In fact, in prior work it has been shown that significantly increased thermostability of ancestral β-lactamases identified through ASR is coupled with increased conformational flexibility,^58,112^ which in turn controls the evolution of substrate specificity.^58,113^ This said, comparative RMSF analysis across all four protein chains reveals a pronounced increase in flexibility in the region spanning residues 190-225 in all designs (**Figures 3, S3** and **S4**).

**Figure 3.**
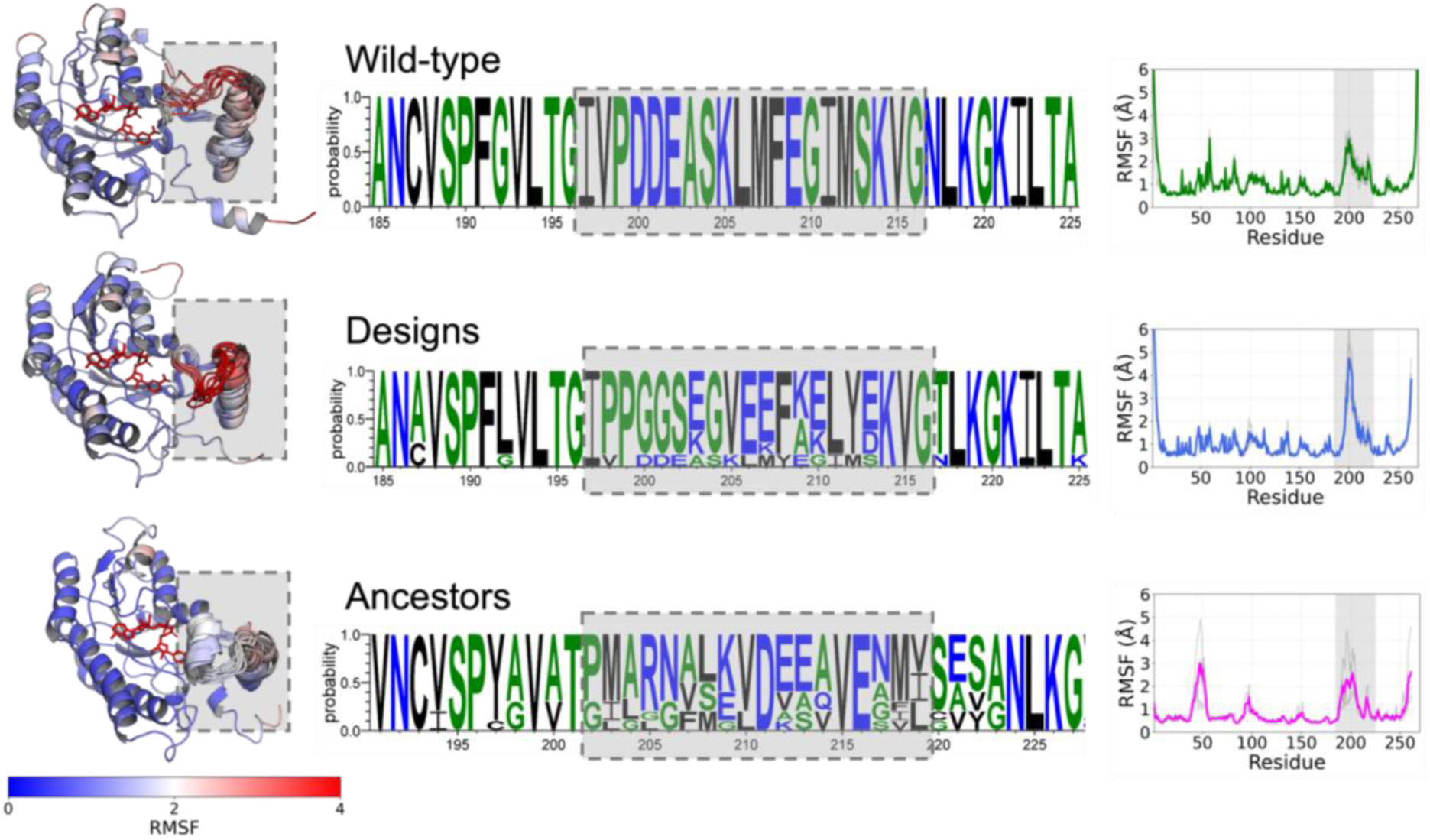
Cartoon representations show ensembles of sampled conformations for residues 185–225, extracted at 50 ns intervals from the last 400 ns of 3 x 500 ns MD simulations of each system, highlighting increased mobility in the wild-type, design 01-080, and ancestor N5. RMSF values are mapped onto the structures using a color scale from blue (low flexibility) to red (high flexibility), corresponding to values between 0 and 4 Å. Detailed RMSF profiles are also shown for wild-type (green), designed variants (blue), and ancestral proteins (magenta). Shown here is also the sequence variability within this mobile region (residues 185-225), calculated using multiple sequence alignment generated from MUSCLE^114^ for the ancestral sequences and illustrated as a WebLogo^115^ comparing wild-type *Sr*BDH1, the FAMPNN^64^ designs, and the ancestors. C_α_-atom root-mean-square fluctuation (RMSF, Å) analysis was performed to compare the scaffold flexibility of wild-type *Sr*BDH1, the FAMPNN^64^ designs, and the ancestral variants from our prior work. The sequence diversification in the highlighted region correlates with altered local flexibility, where designed variants show increased mobility compared to the wild-type and ancestral systems, that may underlie their reduced catalytic activity.

Sequence conservation analysis (**Figure 3**) indicates that residues 200–210 represent one of the most variable segments among the FAMPNN^64^ designs, and that in particular, there is enrichment in glycine residues (which are known to promote backbone flexibility) at positions 200, 201, and 204 providing a higher RMSF in this region (**Figure S3** and **S4**). In contrast, this enrichment is not present at all on the ancestors, and some of them present lower RMSF than the wild-type. This localized increase in conformational dynamics may compromise the precise positioning of structural elements required for efficient catalysis and could therefore impact the enzymatic activity of these variables. The agnostic identification of this loop from our simulations is particularly interesting, given that this loop/helix region has been suggested to play an important role in substrates recognition in other short-chain dehydrogenases (see *e.g.*, refs. ^116–119^). Further, as a point of comparison, we note that in prior work engineering *de novo* Kemp eliminase activity into ASR predicted ancestral β-lactamases, we analogously observed displacement of a key helix as being was essential for introduction of the *de novo* activity.^112^ Further, in these systems, both the *de* novo activity and the significantly increased thermostability of ASR predictions was associated with increased helix and overall scaffold flexibility.^112^

### Experimental Characterization and Validation

Eight redesigns of *Sr*BDH1 were chosen for biochemical characterization based on their predicted change in Gibbs free energy of folding (ΔΔ*G*, **Table S1**). The subset was derived from two design strategies based on residue conservation cutoffs; Strategy 01 (cutoff <4) and Strategy 02 (cutoff <5). The less conservative approach used in Strategy 01 includes redesigns 01-037, 01-080, 01-099 and 01-173, while Strategy 02 includes redesigns 02-037, 02-389, 02-911 and 02-939. With the exception of design 01-173 (**Figure S6**), all redesigns exhibited soluble expression in *Escherichia coli* BL21 (DE3) and were purified (**Figure S7**). In previous work, most BDH from different plants and their reconstructed ancestors could be produced in soluble in *E. coli*. However, a few were found exclusively in the insoluble fraction.^52,55^ While the two redesigns 01-037 and 02-939 show 38% and 64% of the protein yield observed in *Sr*BDH1 of 98 mg/L, all other designs display expression yields between 85-90% of *Sr*BDH1 (**Table S2**). This compares favorably to the protein yield of the thermostabilized ancestor N5 of 18%.^52^

The catalytic performance of the redesigned variants was evaluated in the oxidation of the pure enantiomers (+)- and (–)-borneol, and racemic isoborneol. We observe that Strategy 01 redesigns consistently have about 10-fold higher activity than Strategy 02 redesigns towards all substrates investigated (**Table 1**). Further, Strategy 02 redesigns generally display an up to 10-fold reduction in oxidative rate towards (+)-borneol compared to Strategy 01 redesigns. This suggests that the more conservative redesign approach destabilized the overall catalytic function, leading to a loss in activity. Meanwhile, redesigns of the less conservative Strategy 01 retained 40-65% of the wild-type activity towards (+)-borneol. Notably, and similarly to previously reported ancestors of BDHs,^52^ all redesigned variants show stereopreference for the oxidation of (+)-isoborneol.

**Table 1.**
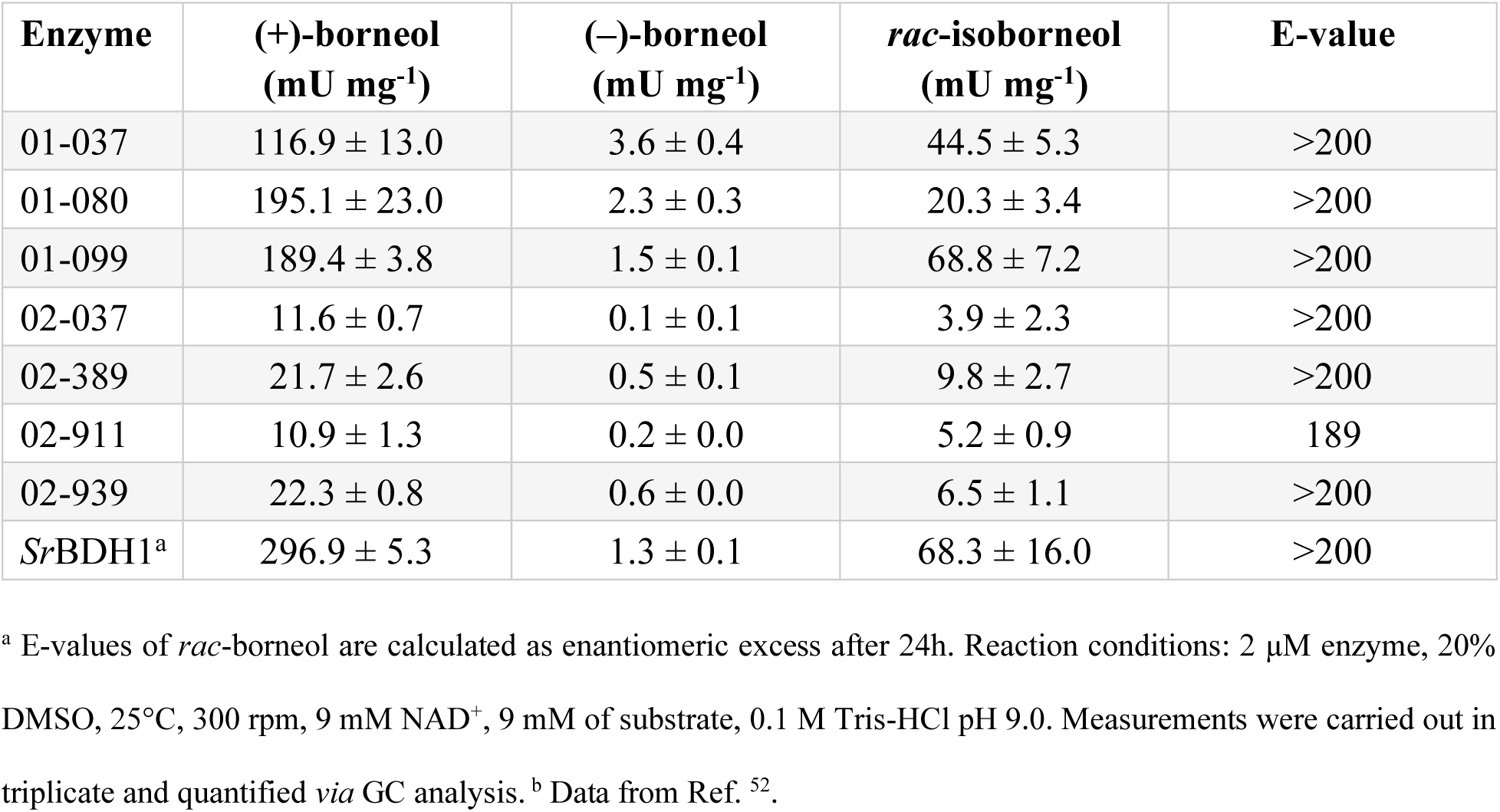
Oxidative rates for (+)-borneol, (-)-borneol, and *rac*-isoborneol given in [mU/mg] catalyzed by the purified redesigns.^a^

The investigation of the enantioselectivity (E-value) in the kinetic resolution of *rac*-borneol confirmed that all redesigns retained exceptional enantioselectivity (E>200). This is a difference to the reconstructed ancestors, where the thermostabilized ancestor N5 has one of the highest (–)-borneol conversion rates, whereas other ancestors with conserved selectivity did not have higher stability than the extant enzyme. The redesign 02-911 had slightly lower selectivity (E=189), which is still sufficiently high for synthetic application. While all redesigns retained the very high E-value, a moderate variation of the relative activities between borneol and isoborneol was observed. 01-080 displays the highest activity toward (+)-borneol, while showing a significant reduction in activity toward *rac*-isoborneol, whereas 01-099 has even lower activity towards (+)-borneol, but converts *rac*-isoborneol faster than 01-080.

Comparison of the designed variants to the ancestral and wild-type variants (**Table 2**) indicates that the redesigned variants show similar *T*_m_ distributions to the thermostable ancestral variants. Within the subset of ancestral proteins, N5 is the most stable ancestor with a *T*_m_ of 77°C. However, N5 displays an E-value of 13 for *rac*-borneol, while the redesign 01-080 retains an E-value of >200. Circular dichroism (CD) spectroscopy was used to cross-validate the DSF data collected for the redesign 01-080 and the wild-type *Sr*BDH1. CD analysis estimated higher denaturation points for both *Sr*BDH1 (71°C) and 01-080 (81°C), while retaining the same difference Δ*T*_m_ of +10°C in design 01-080. Additionally, cooling of the denatured sample back to room temperature, following the initial temperature scan revealed that 01-080 has the capacity for refolding in contrast to *Sr*BDH1 (**Figure S8**).

**Table 2.** Unfolding temperatures (*T*_m_) of the purified redesigned variants, wild-type *Sr*BDH1 and ancestral BDHs, as well as corresponding thermal stability predictions.^a^

| Enzyme | Unfolding Temperature [°C] | SCMTPP Propensity Score | ThermoProt HMM Score | TmProt Score |
| --- | --- | --- | --- | --- |
| *SrBDH1* | 66 | 398.26 | 0.0779 | 48.1 |
| *Designed Variants* |  |  |  |  |
| 01-037 | 67 | 429.72 | 0.137 | 47.7 |
| 01-080 | 76 | 435.15 | 0.121 | 47.4 |
| 01-099 | 70 | 430.91 | 0.091 | 49.4 |
| 02-037 | 69 | 429.66 | 0.136 | 50.8 |
| 02-389 | 68 | 422.97 | 0.141 | 50.8 |
| 02-911 | 65 | 428.54 | 0.163 | 49.9 |
| 02-939 | 68 | 423.69 | 0.076 | 49.6 |
| *Ancestral Variants* |  |  |  |  |
| N4 | 71 | 404.64 | 0.051 | 47.3 |
| N5 | 77 | 409.88 | 0.068 | 47.3 |
| N6 | 66 | 406.86 | 0.052 | 46.8 |
| N7 | 55 | 394.61 | 0.035 | 47.4 |
| N30 | 56 | 404.48 | 0.040 | 48.9 |
| N32 | 61 | 405.07 | 0.126 | 47.3 |
| N39 | 62 | 403.30 | 0.103 | 46.6 |
| N74 | 61 | 403.55 | 0.116 | 47.1 |

<sup>a</sup> Unfolding temperatures for the designed variants were measured *via* differential scanning fluorimetry. Unfolding temperatures for the ancestral variants from ref. <sup>52</sup> were measured *via* CD spectroscopy. Shown here for comparison are also propensity scores calculated using the SCMTPP webserver,<sup>84</sup> ThermoProt Hidden Markov Model (HMM) scores,<sup>52</sup> and predicted melting temperatures, $T_m$ , using TmProt.<sup>120</sup>

The high catalytic activity, thermal stability, and outstanding stereoselectivity of 01-080 thus position it as the best performing variant of either the ancestral or redesigned variants. As such further characterization of the variant was conducted. The apparent Michaelis-Menten constant (*K*_app_) was determined. The redesign displayed a *K*_app_ of 13.7 ± 5.3 mM for (+)-borneol, representing a 6.8-fold increase compared to the wild-type *K*_M_ of 2.02 ± 0.18 mM as reported by Chánique *et. al.*, in 2021 (**Figure S9**).^54^ Due to the solubility limits of the substrate borneol (4mM in water, increased to 10 mM in aqueous solution with 20% DMSO addition) and thus missing saturation of the enzyme, only the apparent Michaelis-Menten constant could be determined. As wild-type SDRs are known to exist as either di- or tetramers,^121^ size-exclusion chromatography (SEC) was performed to validate the quaternary structure of 01-080 in comparison to *Sr*BDH1. The analysis confirmed that the redesigned 01-080 maintains the native tetrameric structure of *Sr*BDH1 without forming higher-order aggregates (**Figure S10**).

Following from this, computational prediction of thermal stability from sequence remains a challenging problem. To this end, we have compared a range of currently available tools for thermostability prediction from sequence, to assess whether they are able to correctly reproduce experimental trends. Impressively, the scoring-card method SCMTPP propensity scores used in the design process (**Figure 1** and **Table 1**) give a significant large positive relationship between experimental and predicted values (Spearman *ρ =* 0.6514, two-tailed *P*-value = 0.0063, see **Supplementary Table**), with propensity scores of 398.26 for the wild-type *Sr*BDH1, 428.7 ± 4.2 for the designed variants, and 404.0 ± 4.4 for the ancestral variants. This qualitatively follows experimental data. In contrast, when comparing to more recent machine learning based approaches (*e.g.*, the Hidden Markov Model (HMM) scores from ThermoProt^52^ or the unfolding temperatures from TmProt,^120^ implemented into EnzymeMiner 2.0^122^), in both cases we do not get significant correlation between predictions and experimental data, and both tools consistently predict all sequences to be moderately stable or not thermostable. For comparison, we also tested whether an older tool, ThermoPred,^123^ which uses a support vector machine based method in order to predict whether a protein sequence corresponds to a thermophilic protein or not, could correctly identify thermophilic proteins in our set, and this tool gave much higher thermophilic probability to the designed sequences compared to the other sequences (see the **Supplementary Table**). The inferior performance of the more modern machine-learning based tools on the designed variants is likely due to dependence on training set, and poor performance when faced with novel sequences. Further, the training sets will be dominated by data on natural and extant sequences, whereas here we are dealing with artificial tetrameric scaffolds. We note that, obviously, two ML-based methods is not exhaustive, and this data was just shown for testing and illustrative comparison.

Finally, while the *T*_m_-value is a good indicator for protein stability, determination of activity losses during incubation under process-relevant conditions is decisive for the catalytic performance. In view of the poor solubility of dibornane-type monoterpenols, conducting the biotransformation at elevated temperatures is desirable. The half-life time (*τ*_(½)_) of 01-080 and *Sr*BDH1 was determined at 50°C, 65°C and 70°C (**Figure S11**). At 70°C, both variants rapidly denatured, with catalytic activity completely lost within 5 min of incubation. In contrast, at 50°C the enzymes were both remarkably stable, complicating reliable determination of the *τ*_(½)_-value. Within a 3-day window of incubation at 50°C, 70% of the activity of 01-080 and 48% in *Sr*BDH1 remained. On day 4, precipitation of both enzymes lead to the termination of the experiment. The most pronounced difference was observed at 65°C. Design 01-080 displayed a *τ*_(½)_ of 4.23 minutes, and retained residual activity of 14% after 30 minutes of incubation, whereas the wild-type *Sr*BDH1 was completely deactivated within the first 5 minutes. The higher residual activity of 01-080 at day 3 of incubation at 50°C, paired with the retention of activity at 65°C highlights a successful stabilization of the redesign in comparison to the wild-type. Furthermore, activity assays conducted at 45°C and 55°C showed a constant activity ratio between the two enzymes (**Table S3**), confirming that while *Sr*BDH1 is functional at elevated temperatures, 01-080 provides a wider operational window for high-temperature biocatalysis.

### Modelling Stereoselectivity in the Most Stable Design (Design 01-080)

Following the strategy established in our previous work,^52^ we assessed putative stereoselectivity toward (+)- and (-)-borneol using short TorchANI-Amber ML/MM simulations,^124–126^ in which borneol was treated at the machine learning level while the protein environment was described classically. Substrate stability in a catalytically active conformation in the active site was assessed by monitoring two key catalytic distances, d_1_, which corresponds to the hydride-transfer geometry between the C_4_ atom of the NAD⁺ nicotinamide ring and the oxidizable carbon of borneol, and d_2_, which describes the hydride transfer between the borneol hydroxyl group and the catalytic tyrosine (**Figure 4**). In our prior work, these distances provided clear discrimination between the different borneol enantiomers in simulations of wild-type and ancestral *Sr*BDH1 variants, with good agreement with experimental selectivity trends.^52^

**Figure 4.**
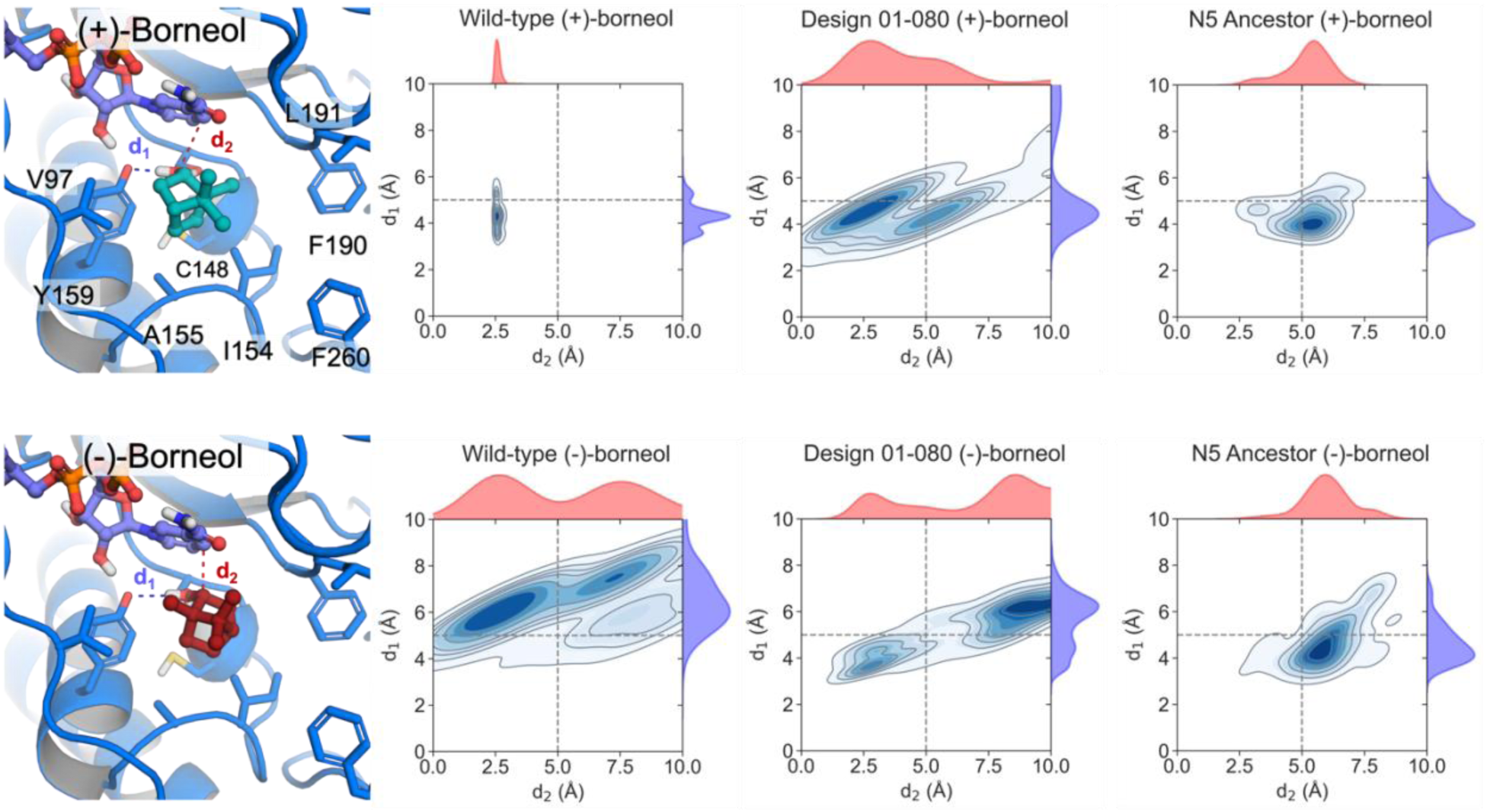
Kernel density estimates (KDEs) of the distribution of key catalytic distances between (+)- and (-)-borneol and wild-type and *de novo* designed *Sr*BDH1, during 3 x 10 ns TorchANI-Amber^126^ ML/MM simulations. Distance d_1_ (blue) corresponds to the interaction between the catalytic tyrosine and the substrate, while d_2_ (red) represents the hydride-transfer donor-acceptor distance. Numbers refer to FAMPNN designs;^64^ for the corresponding sequences see the section “**Wild-type, Designed 01-080 and Reconstructed Sequence N5**” in the **Supporting Information**.

For each variant/enantiomer pair, 10 ns ML/MM simulations were sufficient to stabilize productive binding modes and resolve dynamic differences between (+)- and (–)-borneol. Analysis of the resulting d_1_/d_2_ distance distributions and corresponding kernel density plots (**Figure 4**) reveals that our most stable design, 01-080, selectively maintains catalytically competent geometries for (+)-borneol, while failing to do so for the (–)-borneol (see also **Table S3**). In contrast, while ancestor N6 still shows high activity with (+)-borneol, it is the most active ASR variant on (–)-borneol.^52^ These results indicate a clear preference for (+)-borneol in these designs and suggest that in these *de novo* designs stereoselectivity arises from differential dynamic stabilization of reactive conformations rather than from static binding geometries alone.

## Conclusions

Protein stabilization is an important (but challenging) “Holy Grail” problem for biocatalysis,^1,2^ in particular if one wants to achieve this without compromising activity or selectivity.^3–15^ There has recently been an explosion of approaches for protein stability design (*e.g.*, refs. ^39,43–48,50,51,127^, among others). In the current study, we performed full-atom protein sequence design with sidechain conditioning using FAMPNN,^64^ to engineer thermostable variants of the borneol dehydrogenase from *Salvia rosmarinus* (*Sr*BDH1^54,70^). FAMPNN has previously been suggested to be a powerful tool for design of thermostable and active enzymes.^64^

Multi-objective optimization (in this case enhancing thermostabilty while also retaining enantioselectivity) is challenging in general, and particularly so in the BDH enzyme family, for a number of reasons. (1) This family includes both selective and unselective enzymes towards their natural chiral substrate (1-borneol);^52^ however, unselective enzymes dominate, as stereoselective (+)-1-borneol oxidation does not confer an obvious evolutionary advantage to most plants – with the exception of perhaps those, where the (–)-enantiomer is found as constituent of the essential oils.^71,128–131^ (2) Further, from a structural perspective, the composition of the hydrophobic active site pockets of unselective and selective BDH is very similar,^52^ and point mutations in the highly selective *Sr*BDH1 active site did not reduce its selectivity towards (+)-borneol.^54^ Building on this, detailed molecular simulations indicate that BDH selectivity is driven by dynamical, not structural considerations,^52^ and thus sequence or structure based approaches alone would struggle to predict or rationalize this selectivity. Finally, (3) characterization of ancestral BDHs obtained from ASR indicate that selectivity is lost in ancestors of the highly selective *Sr*BDH1.^52^ While it would be possible to recreate selectivity with first-shell mutations, as discussed, for instance, ref. ^66^, it would be necessary to do so for each unselective variant. It can be argued that similar considerations apply for biocatalysts in which higher selectivity or another catalytic property was introduced by enzyme engineering. It is highly unlikely that reconstructed ancestors of these sequences retain these improved properties, and the re-introduction into the ancestors by mutagenesis could be at best tedious and might not even work (see also refs. ^132,133^ for examples of ASR impacting selectivity and other enzyme characteristics).

By avoiding the *Sr*BDH1 active site residues and guiding our FAMPNN^64^ design protocol with residue conservation filtering using ConSurf,^73,74^ we were able to generate *Sr*BDH1 variants with up to 10 °C enhanced thermostability, while retaining selectivity towards (+)-borneol, unlike the ancestral enzymes identified in our prior work.^52^ Detailed sequence and dynamical analysis indicated once again the importance of conformational dynamics expressed through both scaffold flexibility and active site plasticity in retaining this selectivity, and, encouragingly, use of a scoring card method (SCMTPP)^84^ allowed us to predict thermal stability from sequence in good correlation with experimental measurements (see the **Supplementary Table**). This is significant, because as there is an explosion of ML-based tools to predict enzyme thermal stability (*e.g.*, refs. ^84,123,134,135^) kinetic parameters (e.g., refs. ^136–140^) and even specificity and selectivity (e.g., refs. ^141–143^, note these references are a subset of current studies as the field moves so fast), it will become possible to increasingly automate the design process computationally, minimizing the need for experimental involvement apart from validating top-ranking computational hits, making complete computational design^144^ a reality. This provides a transformative leap forward for biocatalysis.

## Materials and Methods

### ConSurf analysis

Residue conservation scores were calculated using the ConSurf webserver^73,74^ following the standard workflow, including homolog retrieval using the HMMER websever,^145^ multiple sequence alignment with MAFFT,^146,147^ and Bayesian estimation of evolutionary conservation (for methodological details, see ^73,74^).

### Chai-1 models

Initial enzyme-ligand complexes were built with Chai-1,^148^ starting from the *Sr*BDH1 FASTA sequence (see the **Supporting Information** for sequence), together with the SMILES representations of NAD^+^ and borneol. Protonation states were assigned for pH 7 using the H++ server,^149,150^ with the catalytic tyrosine constrained to its deprotonated form. Hydrogen atoms for the oxidized NAD^+^ cofactor was introduced with Open Babel.^151^

### MD simulation setup

Starting coordinates were obtained from Chai-1,^148^ as described above. Protein residues were described using the AMBER ff19SB force field.^152^ Force-field parameters for NAD^+^, borneol and the deprotonated tyrosine residue were derived by restrained electrostatic potential (RESP) fitting^153^ at the HF/6-31G(d) level of theory, using Antechamber,^154^ based on gas-phase geometries optimized at the B3LYP/6-31G(d) level of theory using Gaussian 16 Rev. B.01.^155^ All other non-standard parameters were obtained using the General AMBER Force Field (GAFF2).^156^ All systems were assembled using the tleap module of AmberTools25,^157^ and subjected to an initial vacuum energy minimization using the *sander* module of AMBER24.^157^ This was followed by solvation in a truncated-octahedral box of OPC water molecules,^158^ extending at least 13 Å from the protein in all axes.

Each solvated system was then minimized in three successive stages, relaxing first hydrogen atoms, then the solvent, and finally the complete system. Heating was performed from 100 to 300 K over 50 ps of simulation time under NVT conditions, using a Langevin thermostat (collision frequency 1.0 ps⁻¹),^159^ while restraining all solute atoms with harmonic potentials of 40 kcal mol⁻¹ Å⁻² to preserve the initial geometry during equilibration. Then, the systems were switched to a constant pressure (NPT) scheme, using a Berendsen barostat, at 300 K.^160^ Long range interactions were treated using the Particle Mesh Ewald (PME) method,^161^ with an 10 Å cut-off for non-bonded interactions. The SHAKE algorithm^162^ was employed to restrain the hydrogen atoms in water molecules, allowing for a 2 fs step size for all simulations. Equilibration was performed in three replicates per system, using different initial velocities (assigning different random seeds).

Equilibrated structures were then used as starting points for two different sets of simulations: 3 x 500 ns production MD simulations, and 3 x 10 ns hybrid machine learning/molecular mechanics (ML/MM) simulations (one production MD or ML/MM trajectory from each of three equilibration replicates per system). ML/MM simulations were performed using the TorchANI-AMBER^124–126^ interface implemented in AmberTools25.^157^ In these simulations, the borneol ligand was treated as the ML region, and described using a neural-network potential trained on high-level quantum chemical data, enabling an explicit description of polarization and short-range electronic effects at near QM accuracy, while the remainder of the system was modelled at the MM level, using the previously defined parameters for our MD simulations, as in a standard QM/MM calculation. The ML/MM coupling was implemented using electrostatic embedding with a 12Å ML-MM interaction cutoff, and the ANI architecture^1,163,164^ was exploited using an ANI-MBIS-Q model.^165^ Substrate stability was monitored throughout the simulations by tracking two key catalytic distances, namely the interaction between the catalytic tyrosine and the substrate (d_1_), and the hydride-transfer donor-acceptor distance (d_2_) (see **Figure 4**). Prior work demonstrated this protocol can adequately discriminate between borneol enantiomers in ML/MM simulations of a range of BDHs.^52^ Both conventional MD and ML/MM trajectories were analyzed using CPPTRAJ^98^ and MDAnalysis.^99,100^ Data processing, statistical analysis, and visualization were carried out using Matplotlib.^92^

A Zenodo package containing all starting structures, representative input files, representative frames from our simulations in PDB format, simulation checkpoint files and any non-standard parameters can be found at DOI: 10.5281/zenodo.19267529. *Chemicals.* All chemicals were purchased from Sigma Aldrich (Vienna, Austria) or Roth (Karlsruhe, Germany) and used without further purification unless otherwise stated.

### Cloning of the redesigned SrBDH1 variants

Genes of the redesigned *Sr*BDH1 variants (01-037, 01-080, 01-099, 01-173, 02-037, 02-389, 02-911, and 02-939) carrying an N-terminal 6x His-tag and complementary sequences to the vector pET28a(+) were obtained as synthetic genes (Twist Bioscience, USA). pET28a(+) (final conc. 50 ng/µL) was linearized by restriction digest with NcoI HF (1 µL) and XhoI (1 µL) (New England Biolabs, Germany) in Cut Smart buffer (5µL, final reaction volume 50µL) at 37°C for 15 minutes. Enzymes were deactivated at 80°C for 20 minutes. The synthetic genes were assembled and cloned into the vector pET28a(+) via Gibson Assembly following the original protocol.^166^ The reaction was carried out at 50°C, 300 rpm for 45 minutes. Chemo-competent *Escherichia coli* TOP10 cells were transformed with 5 µL of the assembly mix. Isolated plasmids (Wizard Plus SV Minipreps DNA Purification System, Promega, USA) were sequenced by Sanger sequencing (Microsynth AG, Switzerland) and subsequently used for transformation of *E. coli* BL21 (DE3) for expression of the redesigns.

### Investigation of enzyme expression

Expression studies were carried out to investigate the optimal expression conditions of the redesigned variants in *E. coli* BL21 (DE3). Main cultures of *E. coli* BL21 (DE3) pET28a(+) carrying the respective redesigned variant were prepared by inoculating 100 mL of LB medium containing kanamycin (40 mg/L) in baffled shaking flasks, with overnight cultures to a final OD_600_ of 0.2. After the inoculation, the main cultures were incubated at 37°C and 130 rpm until an OD_600_ of 0.8-1 was reached. At OD_600_ of 0.8-1 the protein expression was induced by addition of 1 mM IPTG. Following the induction, the culture was split into 25 ml aliquots and transferred to sterile 50 ml falcon tubes to assess different expression temperatures. Protein expression proceeded overnight at 18, 28 or 37°C and 130 rpm. Before induction, 4 h and 20 h after induction, samples corresponding to an OD_600_ of 1. The samples were centrifuged at 13.000 rpm, 4°C for 10 min (Centrifuge 5415 R, F-45-24-11, Eppendorf, Germany) and the supernatant discarded. The cells were lysed by addition of 100 µl BugBuster Master Mix (Merck Millipore, USA) followed by incubation at 25°C, 350 rpm for 15 min. Samples were subsequently centrifuged at 13,000 rpm, 4°C for 10 min (Centrifuge 5415 R, F-45-24-11, Eppendorf, Germany). The supernatant was collected and the pellet was resuspended in 100 μL of 8M urea for further analysis on SDS-PAGE.

### Cultivation and expression

Main cultures of *E. coli* BL21 (DE3) pET28a(+) carrying the respective redesigned variant were prepared by inoculating 250 mL of LB medium containing kanamycin (40 mg/L) in baffled shaking flasks, with overnight cultures to a final OD_600_ of 0.1. After the inoculation, the main cultures were incubated at 37°C and 130 rpm until an OD_600_ of 0.8-1 was reached. At OD_600_ of 0.8-1 the protein expression was induced by addition of 1mM IPTG. Protein expression proceeded overnight at 28°C, 130 rpm.

### Cell harvest and lysis

Cells were harvested by centrifugation at 4,000 rpm, 4°C for 15 min (Avanti® J-20 XP and JXN-26, rotor JA-10, Beckman coulter, USA). After the centrifugation, the supernatant was discarded and the cell pellet was resuspended in 20 mL of buffer A (100 mM KPi, 500 mM NaCl, 10% glycerol v/v, pH 8.0). The suspension was centrifuged at 4,000 rpm and 4°C for 15 min (Centrifuge 5810R, A-4-62, Eppendorf, Germany). The supernatant was discarded, and the pellets were then either stored at −20°C for later use or resuspended in 20 mL of buffer A for cell lysis. Cell lysis was facilitated by sonication at 4°C (output control: 6, duty cycle 70%, 6 min; Sonifier S-250A, Branson Ultrasonics™, USA). The lysed samples were centrifuged at 13,200 rpm, 4°C for 15 min (Centrifuge 5415 R, F-45-24-11, Eppendorf, Germany) to remove cell debris. The supernatant (cell free extract) was filtered through a 0.2 µm syringe filter (Filtropur S, PES, 0.2 µm, Sarstedt, Germany) and used for protein purification.

### Protein purification

The proteins were purified by affinity chromatography using High Affinity Ni-Charged Resin (GenScript, China) in gravity columns. Columns contained 5 mL of the sepharose resin. The columns were washed with three volumes of ddH_2_O, and adjusted with two volumes of buffer A, before chilling on ice under rotary shaking at 150 rpm (Titramax 1000, Heidolph Instruments, Germany) for 15 min. Next, the filtered cell free extract was incubated with the sepharose under rotary shaking on ice, at 150 rpm for 15 min. The flowthrough was collected, and the columns were washed with nine column volumes of buffer B (100 mM KPi, 500 mM NaCl, 30 mM imidazole, 10% glycerol v/v, pH 8.0). Proteins were eluted by application of three column volumes of buffer C (100 mM KPi, 500 mM NaCl, 500 mM imidazole, 10% glycerol v/v, pH 8.0). Gravity columns were washed with three column volumes of buffer C, and ddH_2_O and then stored in 20% EtOH at 4°C. The elution fractions were desalted by dialysis overnight with Spectra/Por® 1 Standard RC Dry Dialysis kits (MWCO 6-8 kDa, Spectrum Labs, SAD) in buffer A at 4°C under rotary shaking at 150 rpm (Titramax 1000, Heidolph Instruments, Germany). Desalted protein samples were aliquoted and stored at −20°C.

### Protein quantification and SDS-PAGE analysis

Protein concentration was determined spectrophotometrically at 562 nm in a 96-well format, following the Pierce™ BCA Protein Assay Kit protocol (Thermo Fisher Scientific, US) in the Plate Reader Synergy (BioTek Instruments, US). Measurements were performed in triplicates. SDS-PAGE was conducted to analyze the purified fractions. The samples were mixed with NuPage Sample Reducing Agent (10x), NuPage LDS Sample Buffer (4x) (Thermo Fischer Scientific, US) and ddH_2_O to a final concentration of 1 mg/ml. Samples were heated at 90°C for 10 min. Precast SDS gels (NuPAGE 4-12% Bis-Tris Gel, Thermo Fisher Scientific, US) were inserted into a XCell SureLock Mini-Cell (Thermo Fisher Scientific, US), filled with 1x NuPAGE MES SDS running buffer (Thermo Fisher Scientific, US). 5 μl of the protein standard (PageRuler Prestained, Thermo Fisher Scientific, US) and 10 μl of the samples (1 mg/mL) were loaded onto the gel. The gel was developed under a current of 200 V, and 110 mA for 35 min (PowerEase®500, Invitrogen, US). Afterward, the gels were stained by submerging them in (de-)staining solution (4.9% v/v phosphoric acid, 6.5% v/v EtOH, 0.011% w/v Coomassie Blue G-250 and 1% w/v β-cyclodextrin) for 40 min at room temperature under slow shaking on a tabletop horizontal shaker. Further destaining was facilitated by ddH_2_O at room temperature and continued shaking for 1 h. *Size-exclusion chromatography (SEC)*. SEC was carried out on an Äkta pure system (Cytiva, USA) equipped with a Superdex 200 Increase 10/300 GL (Cytiva, USA). The system was equilibrated to buffer A. Protein Standard Mix 15-600 kDA (Sigma Aldrich, USA) was dissolved according to the manufacturers protocoll and applied to the column for calibration, before recording the wildtype *Sr*BDH1 and 01-080. Signals were recorded at 220, 280 and 340 nm.

### Differential scanning fluorimetry

Determination of the unfolding temperatures of redesigned variants and *Sr*BDH1 at reaction conditions was conducted with the ThermoFluor protocol.^167^ A 200x solution of SYPRO^TM^ Orange (Thermo Fisher Scientific, US) was prepared by diluting a 5000x stock solution with reaction buffer (0.1 mM Tris-HCl, pH 9.0). Enzyme dilutions of 5, 10, and 15 µM were prepared by the addition of reaction buffer. 45 µl of diluted sample solution or buffer (blank) was combined with 5 µl of 200x SYPRO^TM^ Orange solution. Denaturation of the proteins was determined by an increase in fluorescence upon protein denaturation over increased temperature (7500 Real-time PCR System, Applied Biosystems, US). After an initial holding temperature of 30 sec at 25°C, a linear gradient up to 95°C with a 0.3% increase was applied. Samples were measured in triplicate. The resulting unfolding curves were plotted as fluorescent signal over temperature after blank subtraction. The unfolding temperature (T_m_) was determined as the inflection point (maximum of the first derivative). To minimize noise and improve peak detection, raw data were smoothed using a Savitzky-Golay filter prior to derivative calculation.

### Circular dichroism (CD) spectroscopy

CD spectroscopy of the ancestral variants was carried out using a Jasco J-1500 CD Spectrometer (USA). Samples were prepared at a concentration of 0.1 mg/ml of enzyme in 10 mM KP_i_, 50 mM Na_2_SO_4_ at pH 8.0. The signal was recorded at 194 or 217 nm during a temperature ramp of 1°C/min from 20°C to 95°C. Unfolding temperatures were calculated using Spectral Manager. CD spectra of the wild-type *Sr*BDH1 and the redesigned variant 01-080 were recorded using a Chirascan™ Circular Dichroism Spectrometer (Applied Photophysics Ltd., UK) at 25°C, between 180-260 nm with a step size of 1 nm, at a protein concentration of 2 mg/mL in buffer A. Spectral scans were carried out in triplicate. Thermal denaturation curves of the wild-type *Sr*BDH1 and 01-080 were recorded using a Chirascan™ Circular Dichroism Spectrometer (Applied Photophysics Ltd., UK), between 190 to 260 nm with a step size of 1 nm. A temperature gradient from 25 °C to 100 °C was applied in ramp mode. Protein refolding was probed by cooling the sample back to 25 °C. The melting temperature (T_m_) was determined from the thermal shift of the denaturation curves at 220 nm. T_m_ was determined as the inflection point (maximum of the first derivative) of the thermal shift. To minimize noise and improve peak detection, raw data were smoothed using a Savitzky-Golay filter prior to derivative calculation.

### Determination of the specific activity

Determination of the oxidation rates towards (+)-borneol, (−)-borneol or *r*-isoborneol was performed with purified enzymes. Reactions containing 0.2, or 9 mM of the respective alcohol (dissolved in DMSO), 9 mM NAD^+^, 20% total DMSO (v/v) in 0.1 M Tris-HCl (pH 9.0) were started by the addition of 2 µM purified enzyme (final concentration). The reactions were carried out at 25°C, 300 rpm. Over time, multiple samples were taken and stopped by 1:1 addition of 1 M NaOH. Substrate and product were extracted via a two-step extraction process with 1:2 MTBE, the organic layers were collected and dried over Na_2_SO_4_ and product formation was quantified via gas chromatography. Experiments were carried out in triplicate. The oxidation rates of the wild-type *Sr*BDH1 and the 01-080 variant at elevated temperatures were recorded similarly at 45 °C and 50 °C incubation temperature and treated as described before.

### Determination of k_app_

Kinetic parameters of 01-080 were obtained by measuring initial velocities in the presence of varying concentrations of (+)-borneol (0.5–9 mM) at a constant 9 mM of NAD^+^ under the conditions given above. Reactions were started by the addition of 20 µM purified enzyme. Kinetic parameters were obtained by fitting the obtained data to the Michaelis-Menten equation with SigmaPlot 14.0 (Systat Software). Experiments were carried out in triplicates.

### Determination of E-value

Determination of the stereoselectivity expressed as the enantiomeric ratio (E-value) was carried out by racemic resolution. Reactions containing 9 mM *r*-borneol (dissolved in DMSO), 9 mM NAD^+^, 20% total DMSO (v/v) in 0.1 M Tris-HCl (pH 9.0) were started by the addition of 2 µM purified enzyme (final concentration). Reactions were carried out at 25°C, 300 rpm and stopped after 24 h, and treated as described above.

### Quantitative GC analysis

Achiral GC analysis of borneol and camphor was carried out on a Shimadzu GC-2030 GC-FID system equipped with a ZB-5 column (30 m length, 0.32 mm inner diameter, 0.25 µM film thickness, Shimadzu, JP). Samples were injected in split mode (9.1) with an injection temperature of 250°C and FID temperature of 320°C. The instrument was programmed to an initial oven temperature of 50°C. After 5 min of holding time, a linear increase of 40°C/min up to 300°C was set, followed by 5 min holding. N_2_ was set to a constant flow of 30 ml/min. Chiral analysis was carried out on a Shimadzu GC-2030 GC-FID system equipped with a Hydrodex βTBDM column (25 m length, 0.25 mm inner diameter, Macherey-Nagel, Germany). Samples were injected in split mode (50:1) at an injection temperature of 250°C and FID temperature of 320°C. The instrument was programmed to an initial oven temperature of 90°C. After 5 min of holding time, a linear increase of 2°C/min up to 150°C was set, followed by a linear increase of 45°C/min up to 200°C and (2 min holding). N_2_ was set to a constant flow of 24 ml/min.

### Half-life time determination

Half-life time of 01-080 and *Sr*BDH1 were determined via spectrophotometric measurements. During the measurement, the increase of NADH was monitored at 340 nm within 30 min. Reactions at t_0_ were performed at RT, with 20 µM purified enzyme, 2 mM (+)-borneol (dissolved in DMSO, 1% v/v DMSO in 0.1 M Tris-HCl, pH 9.0. Reactions were started with the addition of 2 mM NAD^+^ (final conc.). At later timepoints, reaction mixes containing the purified enzyme, DMSO and buffer were incubated at 50, 65 or 70°C. After incubation, the samples were cooled to room temperature and (+)-borneol was added. The reactions were started and monitored as described before.

## Acknowledgments

SCLK is the Georgia Research Alliance – Vasser Wooley Chair in Molecular Design at Georgia Tech. We thank Georgia Tech and the Georgia Research Alliance for support of this project. We further acknowledge the National Academic Infrastructure for Supercomputing in Sweden (NAISS) for awarding this project access to the LUMI supercomputer, owned by the EuroHPC Joint Undertaking, hosted by CSC (Finland) and the LUMI consortium through grant agreement no. NAISS 2024/8-16 and NAISS 2025/8-11. This work used the Hive cluster, which is supported by the National Science Foundation under grant number 1828187. This research was supported in part through research cyberinfrastructure resources and services provided by the Partnership for an Advanced Computing Environment (PACE) at the Georgia Institute of Technology, Atlanta, Georgia, USA. B.D.G. would like to thank Dr. Ignacio Pickering and Prof. Dario Estrin for their support on the TorchANI-Amber ML/MM simulations. J.Z. would like to thank Anna Schürfer for her help with the CD measurements.

## Declaration of Interests

S.C.L.K. is an informal consultant for Dayhoff Labs, a technology company that builds foundational AI models to solve problems at the chemistry/biology interface.

## Supporting Information

Additional computational and experimental analysis, all SrBDH1 sequences of relevance to this work, and detailed sequence-based analysis of physicochemical properties are provided as Supporting Information. A Zenodo data package with necessary data to reproduce our simulations is provided at DOI: 10.5281/zenodo.19267529.

## Notes

https://zenodo.org/records/19267529

