## Supplementary Data for "Full-Atom MPNN Based Redesign of Plant Dehydrogenase Enables Thermostability Enhancement Without Loss of Stereoselectivity"

[#] These authors contributed equally.

### Table of Contents


**SUPPLEMENTARY FIGURES** .......... **S3**

**SUPPLEMENTARY TABLES** .......... **S11**

**WILD-TYPE, DESIGNED AND RECONSTRUCTED SEQUENCES** .......... **S14**

*Sequences & unfolding temperatures of wildtype BDH* .......... *S14*

*Sequences & unfolding temperatures of ProteinMPNN constructs* .......... *S14*

*Sequences & unfolding temperatures of ancestral proteins* .......... *S15*

*Selected Positions* .......... *S16*

**SUPPLEMENTARY REFERENCES** .......... **S18**

### Supplementary Figures

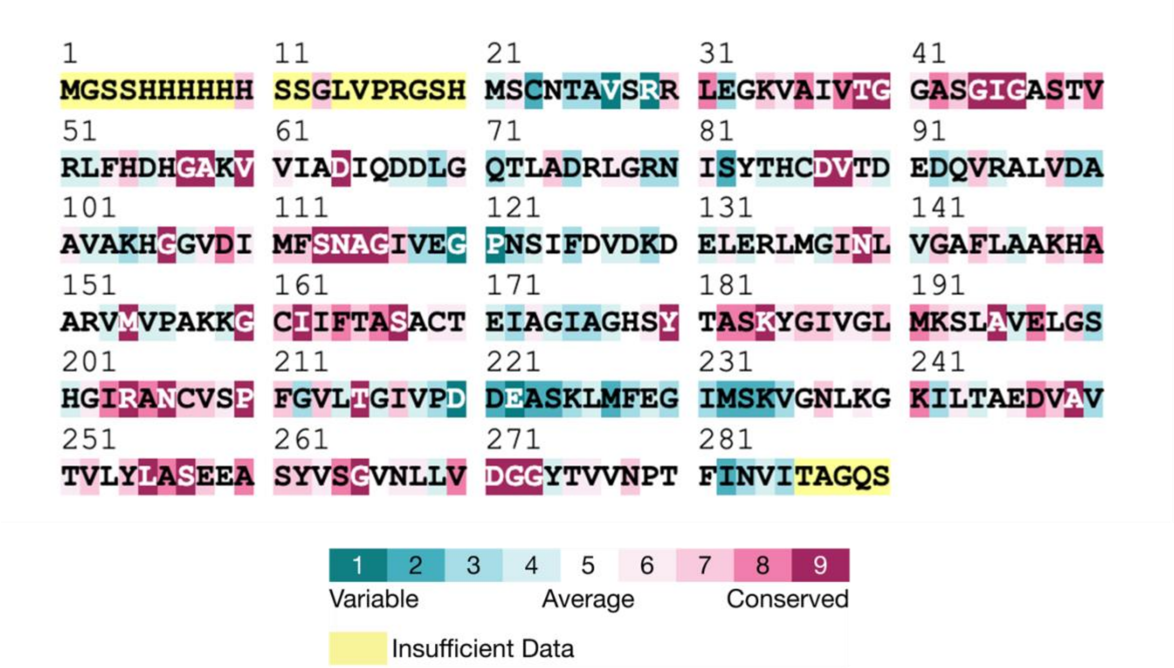


**FIGURE S1.** ConSurf analysis[1,2] of the SrBDH1 sequence (PDB ID: 6ZYZ[3]), showing evolutionary conservation across homologous sequences. Residues are colored according to their conservation scores. Positions with insufficient sequence information for analysis are highlighted in yellow. We note that the positions at the N-terminus highlighted in yellow correspond to the His-tag and linker region of the final construct.

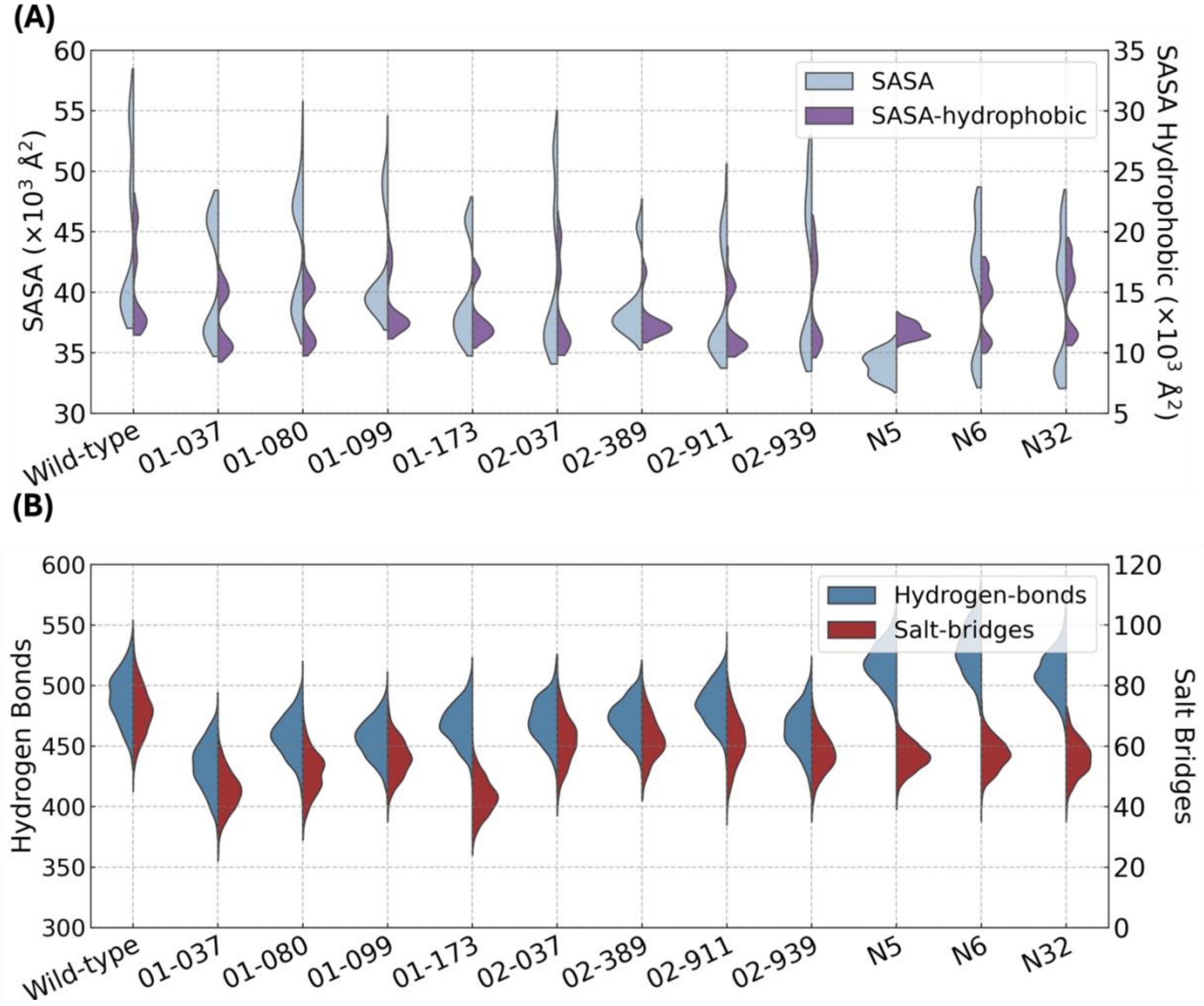


**FIGURE S2.** Violin plots of (**A**) the total and hydrophobic amino acids only solvent-accessible surface (SASA, Å²) and (**B**) the distribution of total hydrogen bonds and salt-bridges, across the last 400 ns of 3 x 500 ns MD trajectories of wild-type *Sr*BDH1 and designed variants, with snapshots taken every ns. Numbers in "xx-yyy" format refer to FAMPNN designs;[4] N5, N6 and N32 correspond to ancestral sequences predicted from our prior work.[5] For the corresponding sequences see the section "**Wild-type, Designed and Reconstructed Sequences**".

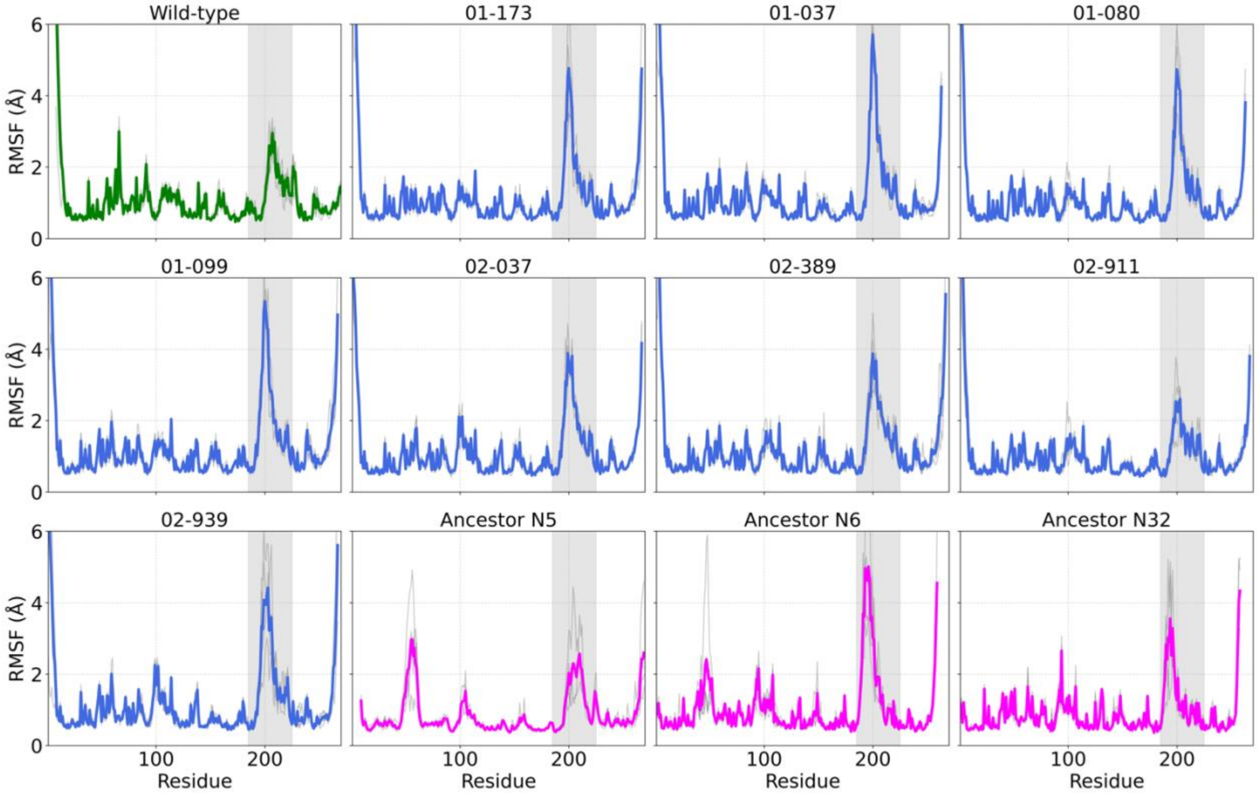


**FIGURE S3.** Root-mean-square fluctuation (RMSF, Å) calculated across the four protein chains of wild-type, designs and ancestors forms of *Sr*BDH1, based on the last 400 ns of 3 x 500 ns molecular dynamics simulations. The individual replicas are colored in grey and the mean values over all replicas are shown in solid color. Region 185 to 225 is highlighted in grey.

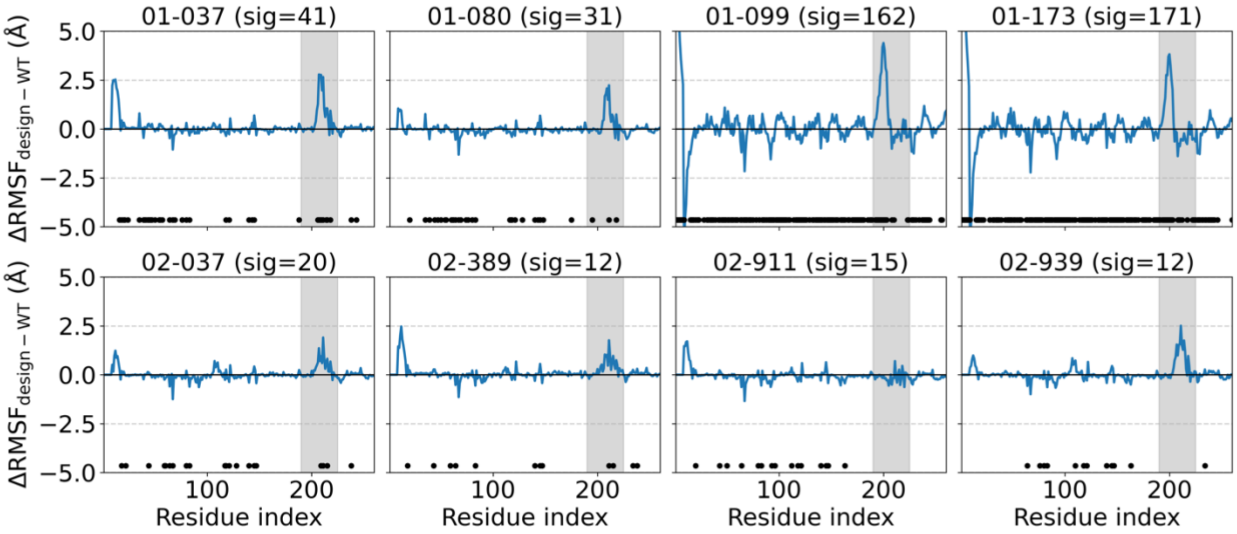


**FIGURE S4.** Differences in root-mean-square fluctuation (ΔRMSF, Å) calculated across the four SrBDH1 protein chains from three independent replicas, based on the last 400 ns of 3 x 500 ns molecular dynamics simulations. ΔRMSF values were obtained as $RMSF_{design} - RMSF_{wildtype}$. Positive values indicate regions that are more flexible in the designs compared to the wild type, while negative values denote regions that are more rigid. The statistical significance of residue-wise ΔRMSF values was assessed using a two-sided Welch t-test,[6] comparing RMSF values obtained from independent replicas of each design against those of the wild-type system. To account for multiple hypothesis testing across residues, *p*-values were adjusted using the Benjamini–Hochberg procedure[7] with a false discovery rate (FDR) of 5%. Residues with adjusted *p*-values below 0.05 are indicated by black dots, and the number of significant residues per system is reported on the panel titles (sig).

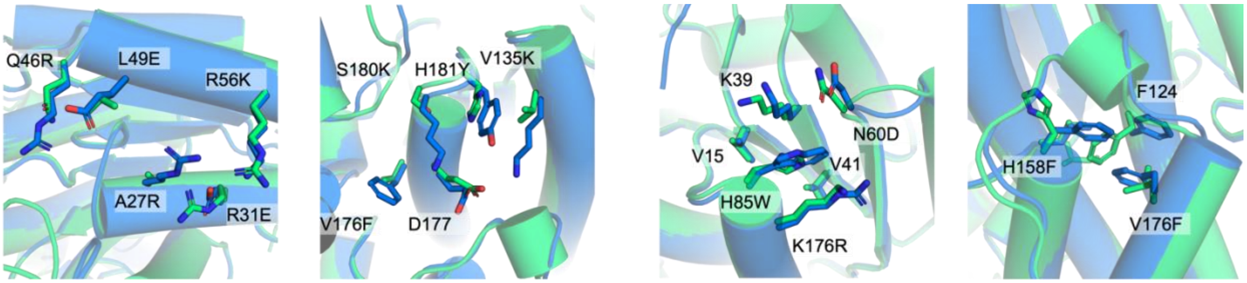


**FIGURE S5.** Cartoon representation comparing wild-type *Sr*BDH1 (green) and design 01-080 (blue), highlighting mutated positions, with newly established interactions shown as sticks.

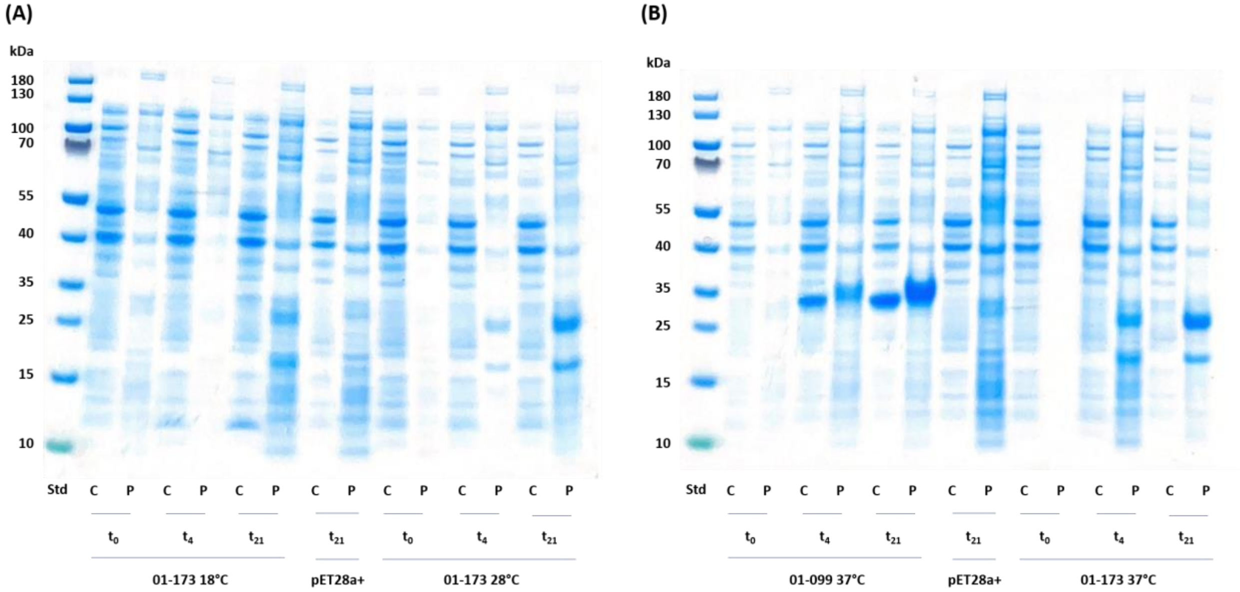


**FIGURE S6.** SDS-PAGE analysis of designs (**A**) 01-173 and (**B**) 01-099 and 01-173, expressed from pET28a(+) in *E. coli* BL21(DE3) at varying temperatures (18°C, 28°C, or 37°C). Samples were collected at three time points (0 h, 4 h, and 21 h), and normalized to an $OD_{600}$ of 1.0. STD = PageRuler Prestained Protein Ladder (Thermo Scientific). C = cell-free extract. P = insoluble cell debris.

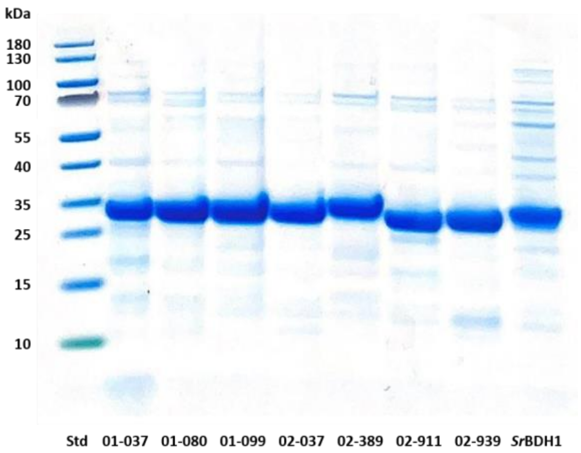


**FIGURE S7.** SDS-PAGE of redesigns, expressed from pET28a(+) in *E. coli* BL21(DE3) at 28°C, and purified via affinity chromatography. Samples are normalized to 1 mg/ml protein loading. STD = PageRuler Prestained Protein Ladder (Thermo Scientific).

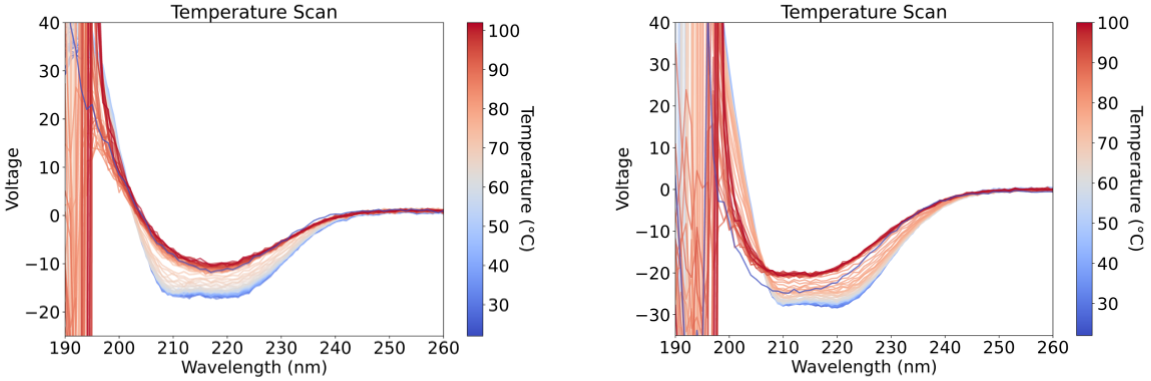


**FIGURE S8.** Temperature scan of (left) *Sr*BDH1 and (right) 01-080 conducted *via* circular dichroism (CD)-spectroscopy, in wavelength ranges from 190-260nm, at 25-100°C (blue to red). After recording the spectrum at 100°C, the sample was cooled to 25°C to probe refolding capacities.

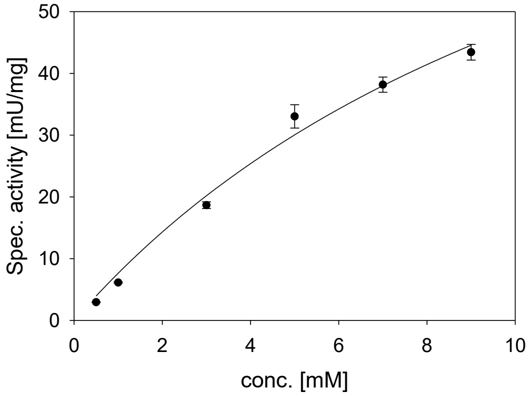


**FIGURE S9.** Measurement of the apparent Michaelis-Menten constant ($K_{app}$) of design 01-080. The specific activities of 01-080 were plotted against (+)-1-borneol concentration and fitted to the Michaelis-Menten regression, visualized in Sigma Plot.[8] Error bars represent the standard deviation of measurements carried out in triplicate, analyzed *via* GC-FID (gas chromatography-flame ionization detector) analysis.

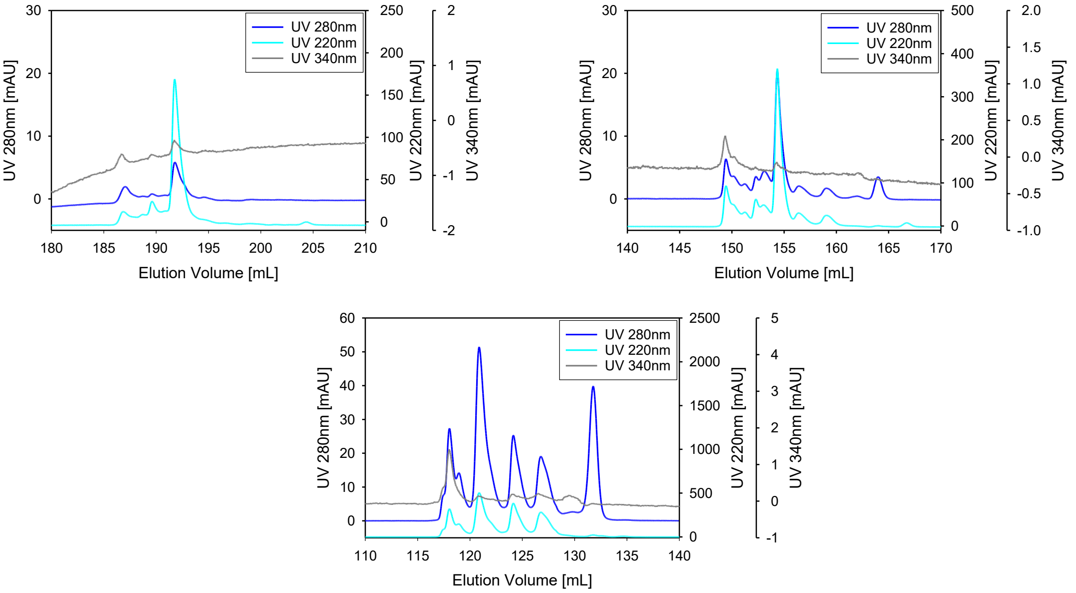


**FIGURE S10.** Size-exclusion chromatography (SEC) carried out on a Superdex 200 Increase 10/300 GL (Cytiva, USA) and monitored at 220nm (red), 280nm (blue) and 340nm (pink). (top left) *Sr*BDH1; (top right) 01-080, (bottom left) Protein Standard Mix 15-600 kDA (Sigma Aldrich, USA).

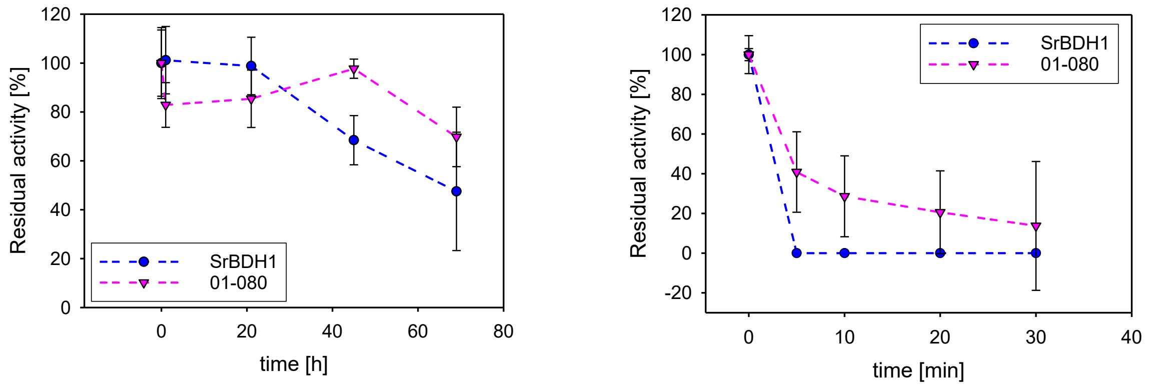


**FIGURE S11.** Determination of the half-life time $\tau_{(½)}$ of design 01-080 (pink triangle) and wild-type *Sr*BDH1 (blue circle) in 1% v/v DMSO in 0.1 M Tris-HCl, pH 9.0. (left) Incubation at 50°C; (right) Incubation at 65°C.

### Supplementary Tables

**Table S1.** Comparison of calculated physico-chemical properties of wild-type SrBDH1, *de novo* designed variants, and selected ancestors.[a]

| System | PyRosetta (REU, ΔΔG) | Chai-1 ipTM | pTM | SASA (Å²) |
|---|---|---|---|---|
| Wild-type | 0 | 0.93 | 0.95 | 44469 ± 6781 |
| 01-173 | -56.41 | 0.93 | 0.95 | 41089 ± 4685 |
| 01-099 | -50.24 | 0.93 | 0.95 | 43310 ± 4824 |
| 01-080 | -48.41 | 0.95 | 0.94 | 41808 ± 4330 |
| 01-037 | -47.9 | 0.93 | 0.95 | 39247 ± 3742 |
| 02-911 | -40.61 | 0.93 | 0.95 | 40825 ± 6344 |
| 02-939 | -36.01 | 0.93 | 0.94 | 38728 ± 2770 |
| 02-037 | -35.4 | 0.93 | 0.94 | 38984 ± 4562 |
| 02-389 | -35.07 | 0.92 | 0.94 | 41641 ± 6438 |
| N5 | - | 0.95 | 0.96 | 34027 ± 1018 |
| N6 | - | 0.95 | 0.96 | 40780 ± 5275 |
| N32 | - | 0.95 | 0.96 | 39351 ± 5223 |
| | **SASA Hydrophobic (Å²)** | **Hydrogen Bonds** | **Hydrophobic contacts** | **Salt Bridges** |
| Wild-type | 15514 ± 3732 | 490 ± 21 | 725 ± 21 | 45 ± 6 |
| 01-173 | 12758 ± 2430 | 434 ± 20 | 772 ± 11 | 45 ± 7 |
| 01-099 | 13322 ± 2455 | 458 ± 18 | 772 ± 44 | 51 ± 7 |
| 01-080 | 13772 ± 2392 | 456 ± 17 | 746 ± 21 | 56 ± 6 |
| 01-037 | 12949 ± 2052 | 468 ± 16 | 767 ± 30 | 43 ± 6 |
| 02-911 | 12383 ± 252 | 471 ± 18 | 778 ± 8 | 62 ± 8 |
| 02-939 | 14453 ± 3703 | 473 ± 16 | 760 ± 39 | 62 ± 7 |
| 02-037 | 13532 ± 3607 | 485 ± 17 | 778 ± 11 | 62 ± 8 |
| 02-389 | 12677 ± 1626 | 464 ± 19 | 775 ± 15 | 57 ± 7 |
| N5 | 11834 ± 616 | 518 ± 17 | 737 ± 4 | 56 ± 5 |
| N6 | 14097 ± 2503 | 529 ± 21 | 708 ± 22 | 57 ± 5 |
| N32 | 14686 ± 2845 | 510 ± 18 | 722 ± 28 | 56 ± 6 |

[a] Shown here are: (1) computational stability estimates calculated using PyRosetta,[9] with ΔΔG shown in Rosetta Energy Units (REU). (2) Interface predicted template modeling (ipTM) and (3) predicted TM-score (pTM) from Chai-

1.[10] (3) Total and hydrophobic-contribution-only solvent accessible surface areas (SASA, Å$^2$) and hydrophobic SASA (Å$^2$). (4) Non-covalent interaction analysis (total hydrogen bonds, hydrophobic contacts and salt bridges). PyRosetta[9] REU scores[11] and Chai-1[10] pTM scores are based on single structure predictions. All other properties are reported as mean ± standard deviation values based on analysis of the last 400 ns of 5 x 500 ns trajectories of each system, with snapshots taken every ns. Numbers refer to FAMPNN[4] designs; for the corresponding sequences see the section "**Wild-type, Designed and Reconstructed Sequences**".

**TABLE S2**. Enzyme yield in mg protein per litre of cultivation media.[a]

| **Enzyme** | **Protein yield (mg $L^{-1}$ of culture)** |
|---|---|
| 01-037 | 38.0 |
| 01-080 | 86.0 |
| 01-099 | 85.0 |
| 01-137 | - |
| 02-037 | 89.0 |
| 02-389 | 83.0 |
| 02-911 | 63.0 |
| 02-939 | 85.0 |
| *Sr*BDH1 | 98.0 |
| N5[b] | 17.8 |
| N6[b] | 45.9 |
| N32[b] | 92.0 |

[a] Expression from pET28a(+) in E. coli BL21(DE3) at 28°C with 1 mM IPTG, overnight. Purified fractions after His-tag affinity chromatography. [b] Data from ref. [5].

**Table S3.** Initial activities of 01-080 and *Sr*BDH1 in mU $mg^{-1}$ determined at 25°C, 45°C and 55°C *via* GC analysis.[a]

| Enzyme | 25°C (mU $mg^{-1}$) | 45°C (mU $mg^{-1}$) | 55°C (mU $mg^{-1}$) |
|---|---|---|---|
| 01-080 | 195.1 ± 23.0 | 355.4 ± 50.8 | 661.3 ± 28.0 |
| *Sr*BDH1 | 296.9 ± 5.3[a] | 473.1 ± 53.5 | 730.0 ± 31.6 |
| *Ratio (%)* | 66 | 54 | 65 |

[a] Ratios of 01-080 oxidative activity over SrBDH1 are given in %. [b] Data from ref. [5].

#### Wild-Type, Designed and Reconstructed Sequences

##### *Sequences & unfolding temperatures of wildtype BDH*

>SrBDH1_Tm_66
MSCNTAVSRRLEGKVAIVTGGASGIGASTVRLFHDHGAKVVIADIQDDLGQTLADRLGR
NISYTHCDVTDEDQVRALVDAAVAKHGGVDIMFSNAGIVEGPNSIFDVDKDELERLMGI
NLVGAFLAAKHAARVMVPAKKGCIIFTASACTEIAGIAGHSYTASKYGIVGLMKSLAVE
LGSHGIRANCVSPFGVLTGIVPDDEASKLMFEGIMSKVGNLKGKILTAEDVAVTVLYLA
SEEASYVSGVNLLVDGGYTVVNPTFINVITAGQS

##### *Sequences & unfolding temperatures of ProteinMPNN constructs*

>01_099_Tm_70
SRLKGKVAVVTGGASGIGASTVRLFHEEGAKVVIADIKEEAGQALAEELGDDVTFVHCD
VTDEEQVKALFDETVERWGGVDIMFSNAGIVEGPNSISDVDKAHFERLMGINLVGAFLT
AKYAAKVMKPQKSGVIIFTASACTEIAGIAGFAYTASKYGVVGLMKELAFELGKYGIRA
NCVSPFLVLTGIPPGGSEGVEKFKELYEKVGTLKGKILTKEDVAKTVLYLASDEASFVSG
VNLLVDGGYTVVNPTFINVVTA

>01_080_Tm_76
GRLEGKVAIVTGGASGIGRSTVELFHEEGAKVVIADIREEEGQALAEKLGDDVTFQHCD
VTDEEQVKALVEATVERWGGVDIMFSNAGIVEGPNSIADVDKADFERLMGINLVGAFL
TAKYAAEVMKPQKSGVIIFTASACTEIAGIAGFAYTASKYGVVGLMKELAFELGKYGIR
ANAVSPFLVLTGIPPGGSKGVEEFAKLYEKVGTLKGKILTADDVAKTVLYLASDEASFV
SGVNLLVDGGYTVVNPTFVNVVNA

>01_037_Tm_67
GRLEGKVAIVTGGASGIGRSTVELFHEEGAKVVIADIREEEGQALAEKLGDDVTFQHCD
VTDEEQVKALVEATVERWGGVDIMFSNAGIVEGPNSIADVDKADFERLMGINLVGAFL
TAKYAAEVMKPQKSGVIIFTASACTEIAGIAGFAYTASKYGVVGLMKELAFELGKYGIR
ANAVSPFLVLTGIPPGGSKGVEEFAKLYEKVGTLKGKILTADDVAKTVLYLASDEASFV
SGVNLLVDGGYTVVNPTFVN

>02_911_Tm_65
GRLEGKVAIVTGGASGIGASTVRLFHEEGAKVVIADIQEEKGKALAEELGDDITFQHCDV
TDEEQVKKLVEDTVEKFGKVDIMFSNAGIVEGPNSISDVDKADFERLMGINLVGAFLTA
KYAAKVMVPQKSGCIIFTASACTEIAGIAGFSYTASKYGIVGLMKSLAFELGKYGIRANA
VSPFLVLTGIPPGGSEGVEEYKELYEKVGTLKGKILTAEDVAKTVLYLASDEASFVSGVN
LLVDGGYTVVNPTFVNVITA

>02_939_Tm_68
GRLEGKVAIVTGGASGIGASTVRLFHEEGAKVVIADIQEEKGQALAEELGENITFQHCDV
TDEEQVKKLVEDTVAKFGGVDIMFSNAGIVEGPNSIFDVKKADFERLMGINLVGAFLTA
KYAAKVMVPQKSGCIIFTASACTEIAGIAGFSYTASKYGIVGLMKSLAFELGKYGIRANA
VSPFLVLTGIPPGGSEGVEEFAELYEKVGTLKGKILTAEDVAKTVLYLASDEASFVSGVN
LLVDGGYTVVNPTFVNVVTA

>02_037_Tm_69
GRLEGKVAIVTGGASGIGASTVRLFHEEGAKVVIADIQEEKGKALAEELGDNITFQHCDV
TDEEQVKALVEDTVKKFGGVDIMFSNAGIVEGPNSIFDVKKEDFERLMGINLVGAFLTA
KYAAKVMVPQKSGCIIFTASACTEIAGIAGFSYTASKYGIVGLMKSLAFELGKYGIRANA
VSPFLVLTGIPPGGSEGVEEFKELYDKVGTLKGKILTAEDVAKTVLYLASDEASFVSGVN
LLVDGGYTVVNPTFINVVTA

>02_389_Tm_68
GRLEGKVAIVTGGASGIGASTVRLFHEEGAKVVIADIQEEKGKALAEELGENITFQLCDV
TDEEQVKKLVEDTVKRFGGVDIMFSNAGIVEGPNSIADVDKADFERLMGINLVGAFLTA
KYAAKVMVPQKSGCIIFTASACTEIAGIAGFSYTASKYGIVGLMKSLAFELGKYGIRANC
VSPFLVLTGIPPGGSKAVEEFKELVEKVGTLKGKILTAEDVAKTVLYLASDEASFVSGVN
LLVDGGYTVVNPTFINVVTA

##### *Sequences & unfolding temperatures of ancestral proteins*

>N4_Tm_71
MASSSSLSSSAKRLEGKVALITGGASGIGECTARLFAKHGAKVVIADIQDDLGQAVCED
LGSESASYVHCDVTIESDVENAVDFAVSKYGKLDIMFNNAGILDPPKPSILDNEKSDFER
VLSVNVTGVFLGMKHAARVMIPARSGSIISTASVASVIGGVASHAYTCSKHAVVGLTKN
VAVELGQYGIRVNCVSPYAVATPMARNFLKLDEEAVENMVSVYANLKGVVLKAEDVA
EAALYLASDEAKYVSGHNLVVDGGFTIVNPSFGMFKQPPNS

>N5_Tm_77
RLEGKVALITGGASGIGESTVRLFVKHGAKVVIADIQDDLGQAVCEDLGSTESVSYVHC
DVTIESDVQNAVDFTVSKYGKLDIMFNNAGILGPPNPSILDNDKSDFERVLSVNVTGVFL
GMKHAARVMIPAKKGSIISTASVASVIGGLGPHAYTASKHAVVGLTKNVAVELGQYGIR
VNCVSPYAVATPMARNALKVDEEAVENMVSASANLKGVVLKAEDVAEAALYLASDE
AKYVSGHNLVVDGGFTSVNPSLGMFR

>N6_Tm_66
MSSSSSSSPPAKRLEGKVAIITGGASGIGESTVRLFVQHGAKVVIADIQDDLGQAICEDLG
STENVSYVHCDVTNESDVQNLVDTTVSKYGKLDIMFNNAGILGRPNSSILDTDKSDFER
VLGVNVTGAFLGAKHAARVMIPAKKGCILFTASVASVIGGLGPHAYTASKHAVVGLTK
NLAVELGQYGIRVNCVSPYAVATPMARNALKVDEEAVENMISESANLKGVVLKAEDV
AEAALYLASDEAKYVSGLNLVVDGGFSTVNPSLTMAMKPPNS

>N7_Tm_55
MSGSSSRAPIAKRLEGKVAIITGGASGIGESTVRLFVQHGAKVVIADIQDDLGQSICKDLG
SDENVSYVHCDVTNDSDVQNLVDTTVSKYGKLDIMFNNAGISGNLNSSILDTDNEDFKR
VFDVNVYGAFLGAKHAARVMIPAKKGCILFTSSVASVISGLGPHAYTASKHAVVGLTK
NLCVELGQYGIRVNCISPYAVATPMLRNAMKVDESAVENMISESANLKGVVLKAEDVA
EAALYLASDESKYVSGLNLVVDGGYSTINQSLTMAMKSLSS

>N30_Tm_56
KRLEGKVAIITGGASGIGASAVRLFLENGAKVVIADIQDDLGQAICDKLGENVSYVHCD
VSNEDDIRNLVDTTVAKYGKLDIMFNNAGILDRPYGSILDTEKSDLERVLGVNVVGSFL

GAKHAARVMVPERKGCILFTASACSVIGGLGTHAYTASKHAVVGLMKNLAAELGQYGI
RVNCVSPYGVVTGMARGVSEVDAEQVESMLSESGNLKGAVLKVEDVAQAALYLASDE
ANYVSGLNLVVDGGFSVVNPSLMMAL

>N32_Tm_61
KRLEGKVAIITGGASGIGASAVRLFWENGAKVVIADIQDDLGQAICDKLGKNVSYIHCD
VSNEDDIRNLVDTTVAKYGKLDIMFNNAGIIDRPYGSILDTEKSDLERVLGVNLVGGFLG
AKHAARVMVPQRKGCILFTASACASIAGLGTHAYTASKHAIVGLMKNLAAELGQHGIR
VNCVSPYGVVTGIGRGVSEVDVAQVEAMLSEVGNLKGAVLKVEDVAQAALYLASDEA
NYVSGLNLVVDGGFSVVNPSMMMAL

>N39_Tm_62
SKRLEGKVAIITGGASGIGASTVQLFHENGAKVVIADIQDDLGQAIANKLGKNVCYIHCD
VSNEDDIINLVDTTVAKYGKLDIMYNNAGIIDRPFGSILDTTKSDLERVLGVNLVGAFLG
AKHAARVMVPQKKGCILFTASACTAIAGLSTHAYAVSKYGIVGLAKNLAAELGQHGIR
VNCVSPYGVVTGIGGVSEVDVAVVEAMLSEVGNLKGQILKAEGVAKAALYLASDEAN
YVSGLNLVVDGGFSVVNPTMMKALNPPES

>N74_Tm_61
MASSSLLSAVAKRLEGKVALITGGASGIGECTARLFSKHGAKVVIADIQDDLGQSVCED
LGSESASFVHCDVTKESDVENAVDFAVSKYGKLDIMFNNAGIVDEPKPSILDNEKSDFER
VLSVNLTGVFLGTKHAARVMIPARSGSIISTASIASVIGGVASHAYTCSKHGVVGLTKNA
AVELGQYGIRVNCVSPYAVATPMARNFLKLDEEAVEGVVSVYANLKGVVLKAEDVAE
AALYLASDEAKYVSGHNLVVDGGFTIVNPSFGMF

##### *Selected Positions*

**Selection of mutational positions.** Residues were then selected directly from the 3D structure based on their ConSurf scores, mutating only positions with low conservation scores, used as designable sites. Two thresholds were applied to define the design space: residues with scores ≤ 4 (Batch 1) and ≤ 5 (Batch 2). These selected positions were provided as mutable sites for FAMPNN, producing 1000 variants per batch.

Batch 1:

1–9, 12, 31–32, 35, 39, 47–49, 51–52, 55, 59–60, 62, 64–66, 70, 72–73, 75, 79–80, 82–85, 87, 98–103, 105, 107–109, 111, 113, 117, 121, 126, 133, 135–136, 139, 151–152, 154–157, 176, 180–181, 192, 194, 196, 198–201, 202–205, 206–209, 210–215, 222, 224, 230, 249, 254, 262–265

Batch 02:

1–9, 12–13, 27, 31–32, 35, 39, 42–43, 45–49, 51–53, 55–56, 59–60, 62–66, 70–73, 75–77, 79–80, 82–85, 87, 98–106, 107–111, 113–114, 116–117, 121, 123, 126–127, 131–133, 135–139, 148,

151–152, 154–158, 161, 176, 178, 180–181, 192, 194, 196, 198–201, 202–205, 206–209, 210–216, 218, 220, 222, 224–225, 230, 233, 239, 246, 249, 254, 257, 259–265
